# Characterization of N distribution in different organs of winter wheat using UAV-based remote sensing

**DOI:** 10.1101/2022.11.02.514839

**Authors:** Falv Wang, Wei Li, Yi Liu, Weilong Qin, Longfei Ma, Yinghua Zhang, Zhencai Sun, Zhimin Wang, Fei Li, Kang Yu

## Abstract

Although unmanned aerial vehicle (UAV) remote sensing is widely used for high-throughput crop monitoring, few attempts have been made to assess nitrogen content (NC) at the organ level and its impact on nitrogen use efficiency (NUE). Also, little is known about the performance of UAV-based image texture features in crop nitrogen and NUE monitoring. In this study, eight flying missions were carried out throughout different stages of winter wheat (from the jointing stage to the stage 25 days after flowering) to acquire multispectral images. Forty-three multispectral vegetation indices (VIs) and forty texture features (TFs) were calculated from images and fed into the partial least squares regression (PLSR) and random forest (RF) regression models for predicting nitrogen-related indicators. Our main purposes were to (1) evaluate the potential of UAV-based images to predict NC in different organs of winter wheat and nitrogen agronomic efficiency (NAE); (2) compare the performances of VIs, TFs and the combination of them for nitrogen monitoring. The results showed that the correlation between different features (VIs and TFs) and NC in different organs varied between the vegetative and reproductive phases. Most of VIs were found to be positively correlated with NC, while most of the TFs were negatively correlated with NC. PLSR latent variables extracted from VIs and TFs explained 80% of the variations in NAE. However, no significant differences were found between VIs and TFs in their performance in predicting NC in different organs. This study demonstrated the promise of applying UAV-based imaging to estimate NC and NAE in different organs of winter wheat.

## 1 Introduction

Higher requirements for crop yield and quality are needed in modern society. Nitrogen (N), as a vital macronutrient, has always been regarded as a key factor in improving crop yield and quality (Wang et al., 2016). In order to ensure high yield, excessive use of N fertilizers in agricultural production have been reported in the North China Plain (NCP) (Cui et al., 2008). Excessive use of N fertilizer causes environmental problems such as soil acidification and water pollution(Ju et al., 2009; Schroder et al., 2011). However, insufficient and inefficient (e.g., wrong time) N fertilizer applications affect the photosynthesis of crops, resulting in reduced crop yield and poor quality (Chlingaryan et al., 2018; Sinclair et al., 2019). Efficient N management for improved N use efficiency (NUE) is critical not only for grain yield and quality but also for environment conservation. Thus, continuous monitoring of crop N status is necessary for the planning of N fertilization measures in the vegetative growth phase and for providing valuable information forecasting yield quality in the reproductive phase (Hank et al., 2019).

Traditional methods for crop N status monitoring based on filed destructive sampling and chemical analysis such as the Kjeldahl technique has the disadvantages of being time-consuming, labor-intensive and costly, limiting the progress in accurate and continuous assessment of crop N status in field (Yao et al., 2015; Onojeghuo et al., 2018). A portable chlorophyll meter was first used for the diagnosis of the leaf N content of rice, and achieved great performance (T. et al., 1986). Subsequently, many studies using portable chlorophyll meters such as SPAD-502 for the monitoring of crop NC have been reported (Errecart et al., 2012; Yuan et al., 2016; Kitonyo et al., 2018). Besides, other handheld crop sensors like GreenSeeker, Crop Circle multispectral active canopy sensors have been developed and applied in the diagnosing of crop N status (Li et al., 2008; Stroppiana et al., 2009; Cao et al., 2013). However, most proximal sensing tools face the challenge of limited throughput. In recent years, the newly emerged UAV remote sensing technology has allowed for high-throughput monitoring and mapping of agricultural ecosystems and has been proven to be convenient and efficient for crop N status monitoring (Kalacska et al., 2015).

With the development of UAV technology, it has been widely used in precision agriculture for its low cost, flexibility and high temporal and spatial resolution (Bendig et al., 2015). Monitoring N status using UAVs has been found successful in different crops in previous studies. For example, (Li et al., 2018c) found it held great potential using UAV-based active sensing for monitoring rice leaf N status. An octocopter UAV was used for capturing multi-angular images to estimate the nitrogen content and accumulation of winter wheat at leaf and plant scale, with the highest accuracy obtained for leaf NC from single-angle images (Lu et al., 2019). There are also many studies about N determination using UAV in other crops such as maize (Maresma et al., 2016), winter oilseed rape (Liu et al., 2018) and sorghum (Li et al., 2018b).

Typically, several methods including statistical regression techniques alongside physically based models are adopted in phenotyping. The physically based models have not been fully examined for crop N monitoring so far though better transferability can be offered (Wang et al., 2015). A few studies proposed modification of radiative transfer models such as the N-PROSPECT (Yang et al., 2015) or N-PROSAIL (Li et al., 2018a) for monitoring crop N status at leaf or canopy scale. However, the models are restricted to few crops and the parameters are complex and not convenient to obtain in agricultural practice (Verrelst et al., 2015; Yang et al., 2015), limiting their use in crop N monitoring. Actually, previous works on N diagnosis in crops predominantly adopted statistical regression techniques. Different spectral features were used to establish parametric or nonparametric linkages with crop physiological and biochemical traits including NC and many other N related indicators. A range of studies has used VIs to construct N estimation empirical regression models and achieve great performance (Song et al., 2016; Tilly and Bareth, 2019). Through the combination of different bands, VIs could be sensitive to the differences (e.g., biomass variation among different stages) in crop phenotypes. (Wang et al., 2012) reported an effective approach of leaf N monitoring using three-band VIs both in wheat and rice. (Zhang et al., 2018) constructed the modified simple ratio index, and found it had a great correlation with wheat NUE. Some published VIs were proved to be well correlated with leaf NC of maize and a new optimized red-edge absorption area index was proposed for the estimation of the vertically integrated leaf NC (Wen et al., 2021). However, crop N monitoring based on single VI could be unreliable due to the limited band information of single VI. With the development of numerous algorithms such as parametric regressions, linear nonparametric regression and nonlinear nonparametric regression, one can make full use of the different bands for crop N monitoring based on VIs (Berger et al., 2020). Texture, as an important characteristic for image classification, has been used in the estimation of forest aboveground biomass (Murray et al., 2010; Kelsey and Neff, 2014). Recently, image texture information have been increasingly used for crop monitoring. (Zheng et al., 2019) found that the using the combination of textural information with spectral information derived from UAV-based images could significantly improve the accuracy for rice biomass estimation compared to the use of spectral information alone. (Yue et al., 2019) has also found similar results in winter wheat biomass monitoring. (Zheng et al., 2020) found that the integration of texture information and VIs could significantly improve all N nutrition parameters estimation using multiple linear regression. However, little is known about the feasibility of using image texture information extracted from UAV images for assessing crop NUE indicators.

It is well known that crop growth is a dynamic process with constant nitrogen turnover. The operation of nitrogen varies in different growth stages and different organs in crops (Ohyama Takuji, 2010). Studies have indicated that different organs could have different effects on the crop spectral features (Li et al., 2015, 2021). However, few investigations under field conditions address the differences of estimated the NC in different organs when using UAV-based multispectral data. Therefore, the main objectives of this study are to (1) evaluate the potential of UAV-based remote sensing images to predict NC in different organs of winter wheat; (2) compare the performance of nitrogen monitoring in winter wheat based on VIs, TFs and the combination of them, in combination with regression algorithms.

## 2 MATERIALS AND METHODS

### 2.1 Study Area and Experimental Design

Field trials were conducted at the Wuqiao Experimental Station of China Agricultural University (37°41′N, 116°37′E) in Hebei Province in the North China Plain (NCP) (Figure 1) within the winter wheat season of 2020 to 2021. NCP belongs to a warm temperate semi-humid continental monsoon climate. The average rainfall, temperature and altitude were about 550 mm, 12.5 °C and 18 m. JiMai22 (*Triticum aestivum L*.), one of the most widely grown winter wheat varieties in NCP was used in this study. It was sowed in October 2020 and harvested in June 2021 with a row spacing of 15 cm and a density of 430 × 10^4^ ha^-1^. The experiment followed a block design and five levels of nitrogen fertilizer treatments were established, including 0 kg N ha^-1^ (N0), 120 kg N ha^-1^ (N1), 180 kg N ha^-1^ (N2), 240 kg N ha^-1^ (N3) and 300 kg N ha^-1^ (N4). 120 kg P_2_O_5_ ha^−1^ and 90 kg K_2_O ha^−1^ were applied to the soil as basal dressings and the rest of the field management followed the local crop production standards throughout the winter wheat season. Besides, three replications were conducted for each treatment, and each plot area was 40 m^2^ (10 m × 4 m).

**Figure 1.**
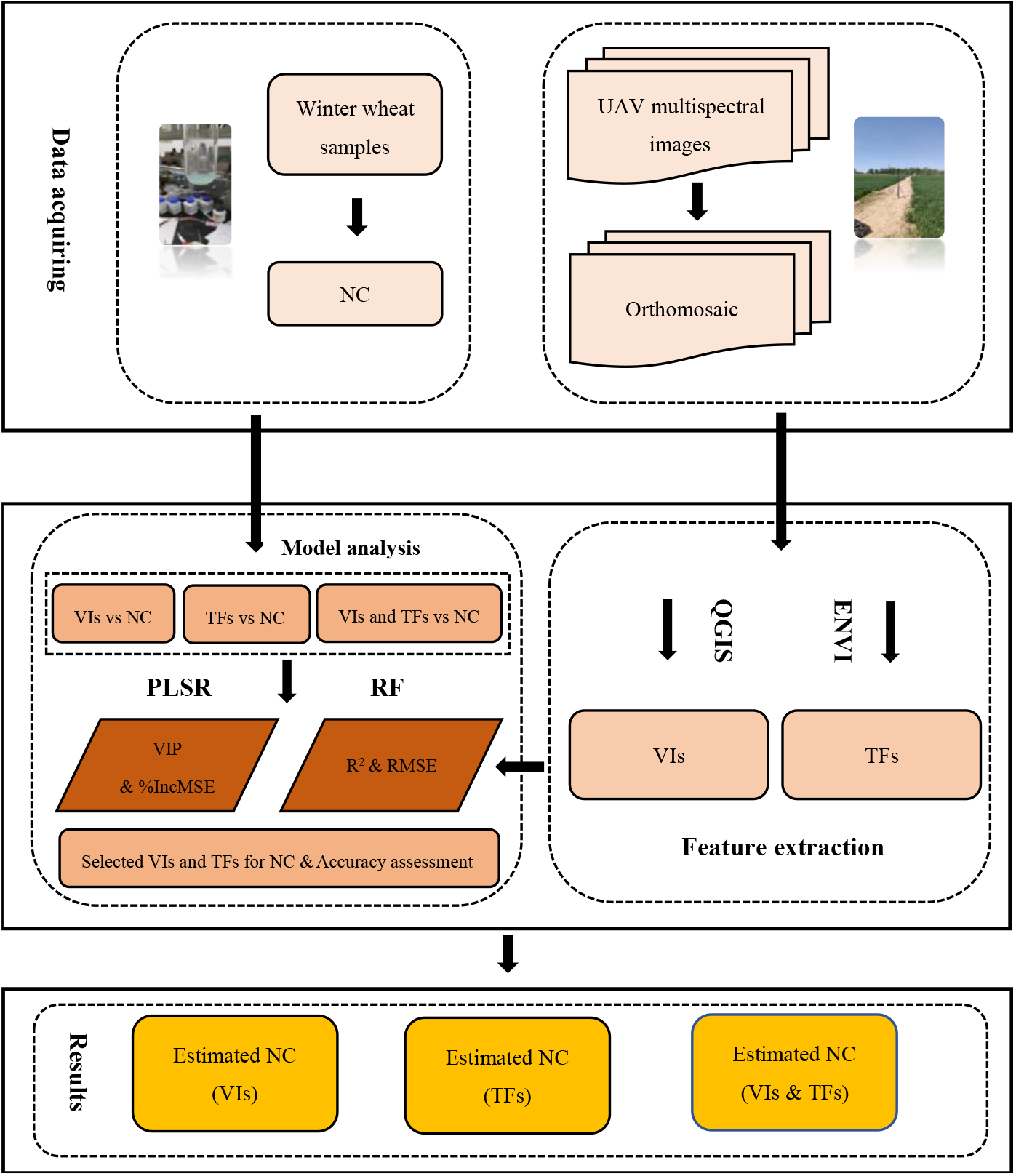
The flowchart of the key steps for data collection and analysis in this study.

### 2.2 Data Collection

#### 2.2.1 Field Sampling and NC Determination

Destructive samplings were performed eight times (dates) during the growth of winter wheat, including three times in the vegetative growth phase and five times in the reproductive growth phase of winter wheat (Table 1).

**Table 1:** Cultivar, treatments and data acquisition schedule.

| Cultivar | N rate (Kg ha<sup>-1</sup>) | UAV Flight Date | Field sampling Date | Growth stage | Zadoks Codes |
| --- | --- | --- | --- | --- | --- |
| JiMai22 | 0 (N0), 80 (N1),<br>120 (N2), 160 (N3), 200 (N4) | 18 April 2021 | 18 April 2021 | Jointing stage (JS) | GS31 |
|  |  | 27 April 2021 | 27 April 2021 | Booting stage (BS) | GS40 |
|  |  | 5 May 2021 | 5 May 2021 | Heading stage (HS) | GS50 |
|  |  | 12 May 2021 | 12 May 2021 | 5 Days after flowering (AF5) | GS70 |
|  |  | 17 May 2021 | 17 May 2021 | 10 Days after flowering (AF10) | GS75 |
|  |  | 22 May 2021 | 22 May 2021 | 15 Days after flowering (AF15) | GS80 |
|  |  | 27 May 2021 | 27 May 2021 | 20 Days after flowering (AF20) | GS85 |
|  |  | 1 June 2021 | 1 June 2021 | 25 Days after flowering (AF25) | GS90 |

Winter wheat plants within an area of 0.06 m^2^ (0.2 m × 0.3 m) of were randomly selected from each plot and transported back to the laboratory immediately. All plants were separated into different organs (leaf, stem, spike and grain). The samples of organs were oven-dried for 30 mins at 105 °C and later at 80 °C to a constant weight. After obtaining the dry matter weight (DMW) of the different organs, dried organ samples were ground to pass through a 1 mm screen and stored in plastic bags for further elemental (N) analysis. At the mature stage of wheat, a 1.8 m^2^ area of wheat plants were randomly harvested from each plot to determine the final yield. The micro-Kjeldahl method (A., 1982) was used to determine the total N concentration of different organs. Equation (1) was used to calculate the plant NC. As one of the indicators for crop NUE, the nitrogen agronomic efficiency (NAE) can be calculated by equation (2).

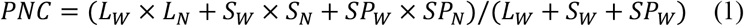

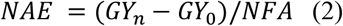

Where L_W_, S_W_, P_W_ were the DMW of leaf, stem and spike, respectively. L_N_, S_N_, SP_N_ were the N concentration of leaf, stem and spike, respectively. And *GY*_*n*_ is the grain yield with N fertilizer application, *GY*_*0*_ is the grain yield without N fertilizer application. *NFA* means the amount of applied N fertilizers (kg/ ha).

#### 2.2.2 UAV Image Acquisition

The acquisition dates of UAV-based images can be found in Table 1. All UAV flight missions were carried out at approximately 10:00 am and 14:00 pm on sunny days. DJI Phantom 4 quadcopter (DJI, Shenzhen, Guangdong, China), which was equipped with a consumer-grade multispectral camera was used in this study. The camera consists of six sensors, including five monochromatic sensors and one Red-Green-Blue (RGB) sensor. The spectral resolution of the monochromatic sensors includes: a blue band with 450 nm center and 16 nm bandwidth, a green band with 560 nm center and 16 nm bandwidth, a red band with 650 nm center and 16 nm bandwidth, a red-edge band with 730 nm and 16 nm bandwidth and a near-infrared band with 840 nm and 26 nm bandwidth. More specific parameters of the UAV and the camera are demonstrated in (Wang et al., 2022a).

Nine ground control points (GCPs) were evenly placed over the field for subsequent image geometry correction. To record the precise coordinate information of GCPs, a D-RTK 2 high-precision GNSS mobile station (DJI, Shenzhen, Guangdong, China) operating at centimeter-level positioning precision with uninterrupted data transmission was used in this experiment. The UAV was flown over the winter wheat field at an altitude of 25 m above the ground level. All flight missions were conducted using the DJI go pro software (DJI, Shenzhen, Guangdong, China), with the heading and side overlaps of 80% and 70%, respectively. All acquired images were saved in TIFF format on the SD card onboard the UAV.

### 2.3 Image processing

#### 2.3.1 Generation of orthophoto maps

We used the Pix4D (Pix4D SA, Lausanne, Switzerland) based on the structure-from-motion (SfM) technique to generate orthophoto images. Following image alignment, matching, mosaicking, sparse point cloud, and dense point cloud constructing, the orthoimages were generated. The ‘Multi-spectral Ag’ template was selected as the processing model for the orthomosaic reflectance images. The coordinates of GCPs were used for orthomosaic georeferencing by manually identifying the points after generating the sparse point cloud. Finally, five georeferenced single-band orthophotos were obtained in each observed stage with the Geo-TIFF format.

#### 2.3.2 Selection and extraction of vegetation index and image texture

Forty-three nitrogen-related VIs (Table S1 in Supplementary Material S1) were screened for further analysis. QGIS (QGIS Version 3.14) was used to calculate the vegetation index maps. We used the function of the “raster calculator” to obtain the VI-maps based on single-band orthophotos generated by Pix4D for each observation stage. Also, eight grey-level co-occurrence matrix (GLCM)-based textures including the mean (Mean), variance (Var), homogeneity (Hom), contrast (Con), dissimilarity (Dis), entropy (Ent), second moment (Sec), and correlation (Cor) (Haralick et al., 1973) were computed using the ENVI software (Exelis Visual Information Solutions, Boulder, Colorado, USA) with the size of moving window of 5 × 5 and in the direction of 45° for all the five single-band orthophotos (Table S2 in Supplementary Material S1). Next, regions of interest (ROIs) were selected for each plot, and the mean values of the VI-maps and texture maps were extracted using the “Zonal Statistic” function in QGIS.

### 2.4 Model development and evaluation

#### 2.4.1 Model calibration

Correlation analysis was performed for the VIs and the nitrogen content of different organs. Meanwhile, to evaluate the performance of the 43 VIs and 40 TFs obtained from the UAV-based images, the Pearson correlations between VIs/TFs and NC of winter wheat were implemented during the vegetative and reproductive growth phase. For further determination optimal combination of multispectral VIs, TFs and regression algorithms for nitrogen prediction, the Partial Least Squares Regression (PLSR) and Random Forest (RF) algorithms were adopted in this study.

Partial least squares regression is one of the most used algorithms to search the basic relationship between two matrices (independent and dependent variables), that is, a latent variable method for modeling the covariance structure in these two vector spaces. It has the advantages of being stable, and suitable for small datasets and can avoid multicollinearity. By conducting the one-sigma algorithm (Wold et al., 2001), the optimal number of latent variables was determined. For the evaluation of the contribution of different VIs to the prediction model, the Variable Importance in Projection (VIP) criterion was introduced (Hastie et al., 2005). In general, variables with a VIP score greater than 1 are considered to be more important to the model. Meanwhile, the larger the VIP value obtained by the variable, the greater the contribution of the variable to the model.

The random forests algorithm was developed by (Breiman and Cutler, 2012) in 2001. As a typical ensemble algorithm, it is composed of multiple unrelated decision trees, and the final output of the model is jointly determined by each decision tree in the forest. It shows a promising capability to avoid overfitting by sampling the predictor space randomly. The number of decision tree (*ntree*) and the input variables per node (*mtry*) are two key hyperparameters that have great impact for the complexity of RF models (Wang et al., 2019). In this study, they were selected based on the root mean square error (RMSE) with the RF algorithm. Besides, as an effective indicator for evaluating the contribution of variables to the model, the percentage increase in mean squared error (%IncMSE) (Farrés et al., 2015) was used in our research. By using the function of ‘rfPermute’ in RF models, the image feature with great importance for the models can be screened out.

All datasets were randomly divided into a training dataset (80%) and a test dataset (20%). The packages “pls” (Mevik and Wehrens, 2007) and “randomForest” (Breiman and Cutler, 2012) were used to construct the prediction models in R programming language in R Studio (R Version 3.6.1).

#### 2.4.2 Model evaluation

The 1:1 line of the estimated and measured nitrogen concentrations were used to assess the fitness of different prediction models. Coefficients of determination (R^2^) and root mean square error (RMSE) were selected to evaluate the performances of the different models. Generally, the higher the R^2^ and the lower the RMSE, the better the precision and accuracy of the models. These statistical indicators were expressed as equations (3) and (4):

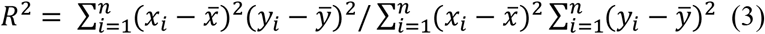

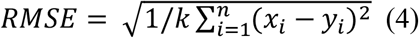

Where *n* is the number of samples, *i* is the ith sample, *x*_*i*_ and *y*_*i*_ stand for the estimated NC values and measured nitrogen concentration values, 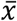 and 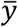 stand for the average estimated NC values and measured NC values, respectively. Figure 1 shows the flowchart of the experiment.

## 3 RESULTS

### 3.1 Measured data from destructive sampling

#### 3.1.1 Descriptive analysis of NC and dry matter weight (DMW)

As shown in Table S3 in Supplementary Material S1, the DMW ranges from 1.22 to 4.18 t/ha with CV of 31.01% in leaf DWM, from 2.72 to 9.87 t/ha with CV of 39.71% in stem DMW, from 0.42 to 2.37 t/ha with CV of 51.53% in spike DMW, and from 3.97 to 15.96 t/ha with CV of 31.71% in plant DMW during the vegetative growth phase. For the reproductive growth phase, leaf-, stem-, spike-, grain- and plant DWM ranges from 0.84 to 3.93 t/ha, 5.14 to 13.80 t/ha, 1.42 to 11.20 t/ha, 0.26 to 8.14 t/ha and 8.26 to 27.08 t/ha, respectively, with CV of 30.76%, 23.44%, 47.86%, 74.41% 24.06%.

Nitrogen content (NC) varies from 2.24% to 4.95%, 0.85% to 1.81%, 1.99% to 4.73%, 1.45% to 3.00% in the leaf, stem, spike and plant, respectively, with CV of 16.84%, 19.48%, 31.01% and 20.86% during the vegetative growth phase. For the reproductive growth phase, the leaf, stem, spike, grain and plant NC varies from 0.91% to 3.64%, 0.29% to 1.31%, 1.41% to 2.60%, 1.62% to 3.06%, and 0.80% to 1.85%, respectively, with CV of 33.41%, 31.10%, 12.00%, 15.04% and 17.78%. It can also be found that the variation of leaf NC and stem NC in the reproductive growth phase was greater than that in the vegetative growth phase (Table S3), which was opposite with the variation trend of spike NC and plant NC in the vegetative and reproductive growth phases.

Figure 2 shows the relationship between the DMW of leaf, stem, spike and plant and the corresponding NC values for the vegetative and reproductive growth phases. Except for the leaf NC, NC in the stem, spike and whole plant decrease as DMW increases due to the dilution effect of N as described in (Lemaire et al., 2008).

**Figure 2.**
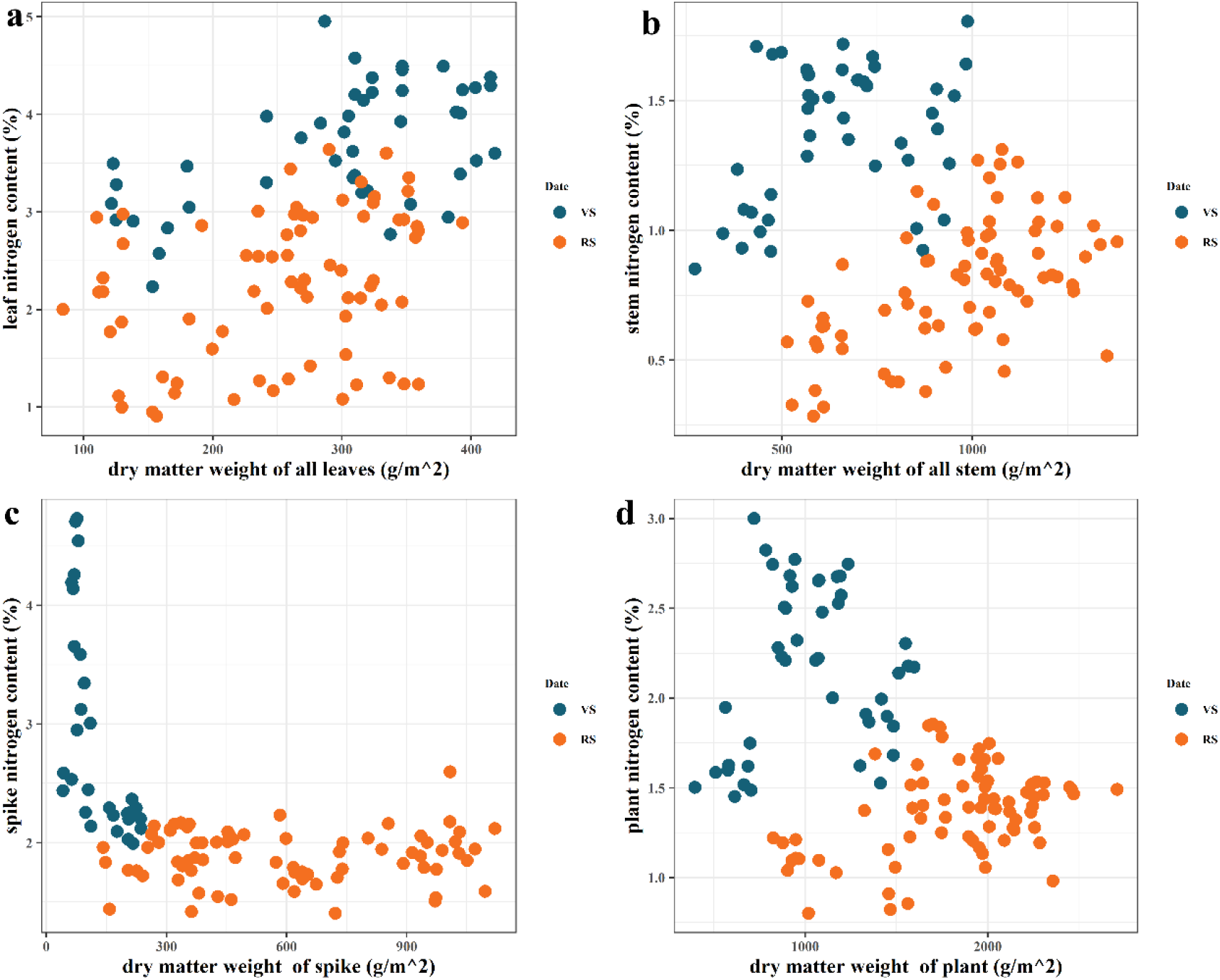
Winter wheat DMW (g/m^2^) vs. winter wheat nitrogen content in the vegetative and reproductive growth phases; (**a**) leaf DMW and leaf NC; (**b**) stem DMW and stem NC; (**c**) spike DMW and spike NC; (**d**) plant DMW and plant NC. VS and RS means the vegetative and reproductive growth phases.

#### 3.1.2 Yield and nitrogen agronomic efficiency (NAE)

Figure 3 depicts the average yield and the corresponding NAE for each N treatment in the experiment. The highest yield was observed in the N3 treatment, whereas the lowest yield was observed in the N0 treatment. NAE decreased significantly along with the increase of N fertilizer inputs.

**Figure 3.**
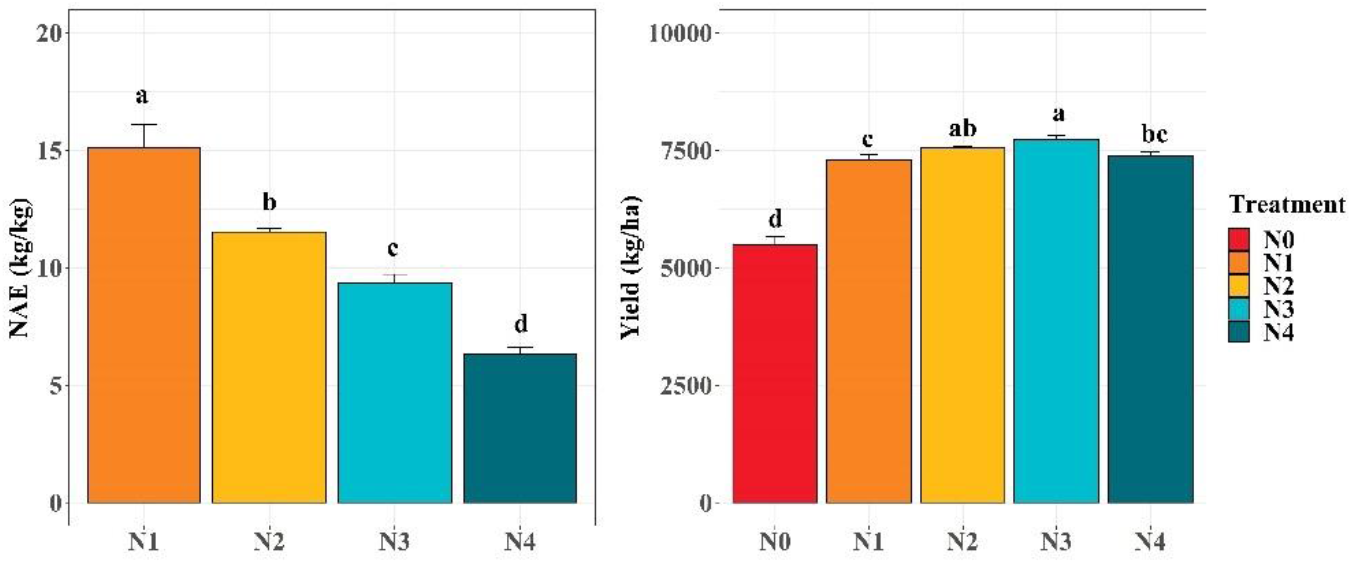
Yield and NAE of each treatment of N. The different small letters indicate significant differences between treatments.

### 3.2 Correlation between image features vs. N-related indicators

Table S4 in Supplementary Material S1 showed the top 5 most relevant VIs and TFs for NC monitoring of winter wheat. In the vegetative growth phase, the RGBVI, MCARI, MCARI2 and RGBVI, were the best correlated VIs for leaf, stem, spike and plant NC, respectively, with r of 0.75, 0.80, 0.60 and 0.75. The Reg_mean (r = -0.85), G_cor (r = -0.84), R_con (r = 0.32) and Reg_mean (r = -0.86) was the best correlated TFs for leaf, stem, spike and plant NC monitoring. In reproductive growth phase, the GOSAVI and R_ho (with r of 0.88 and 0.84), MSR-REG and G_mean (with r of 0.82 and -0.81), DVI-REG and Reg_mean (with r of 0.56 and -0.64), RTVI-CORE and G_mean (with r of 0.71 and -0.58) and CVI and Reg_mean (with r of 0.77 and -0.79) yield the highest r with leaf, stem, spike, grain and plant NC (See detail in Supplementary Material S2).

In general, most of VIs were found to be positively correlated with NC, while most of TFs were negatively correlated with NC. Among all the organs and the whole plant, it was obvious that the correlation between spike NC and image features was the lowest.

Figure 4 shows the absolute value of the r between VIs and NAE in different growth stages. It is clear that the VIs derived from our UAV images can reflect the change of NAE to a certain extent, and the correlation decreases with the winter wheat growth in general.

**Figure 4.**
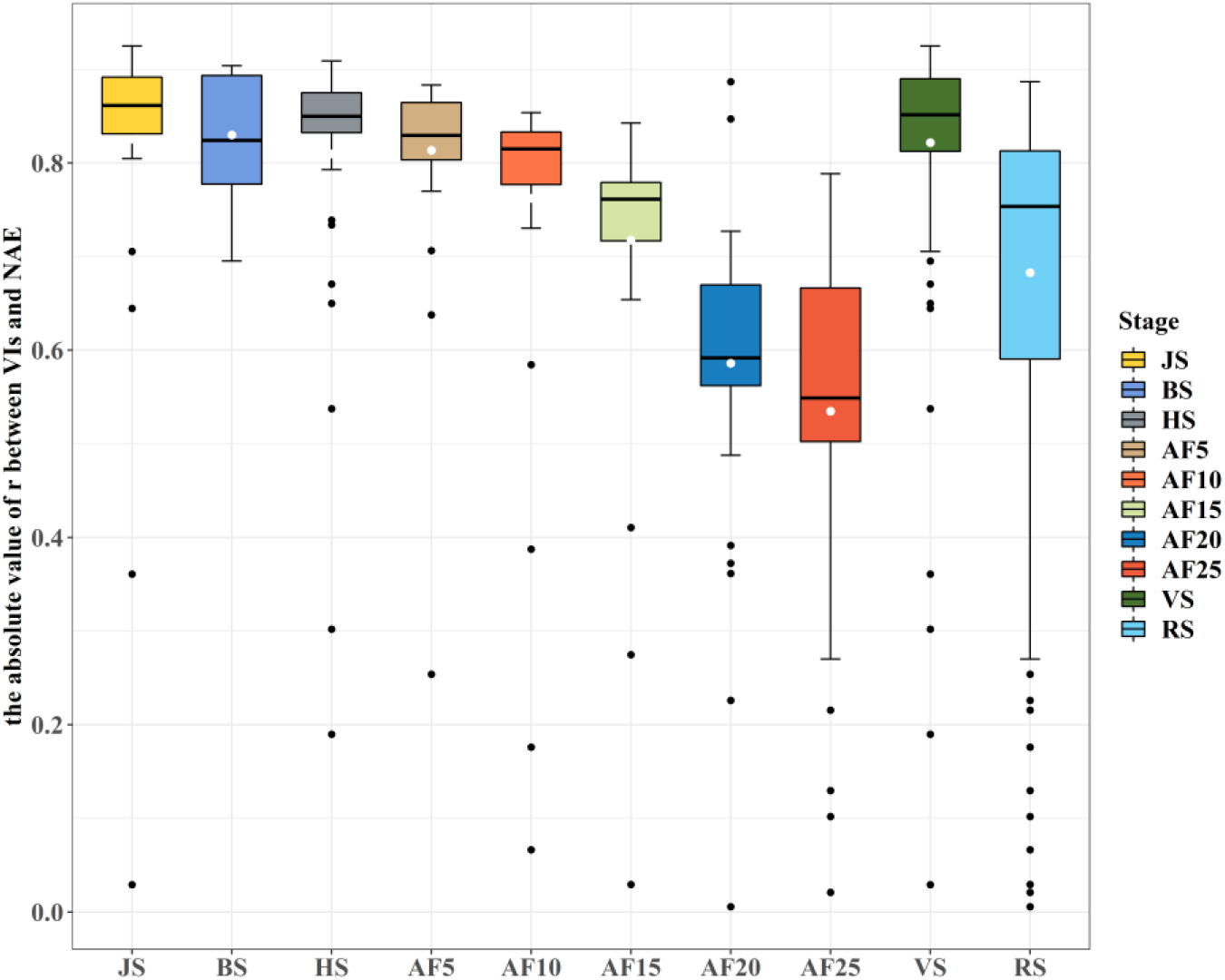
Variation of the absolute value of the r between VIs and NAE in different growth stage. The white dots in each box represent the mean value of the absolute value of the r, and the black dots represent outliers. JS, BS and HS are jointing, booting and heading stage, respectively. And AF5, AF10, AF15, AF20 and AF25 means 5, 10, 15, 20 and 25 days after flowering. VS and RS refer to the vegetative and reproductive growth phases, respectively.

### 3.3 PLSR and RF models using VIs for nitrogen content estimation

As shown in Table 2, during the vegetative growth phase, the PLSR model obtained the highest R^2^ in spike NC estimating both in training and testing sets but the RMSEs were also generally larger than the ones in the PLSR models. For other organs or the whole plant, there were no obvious differences in the estimation during the vegetative growth phase (R^2^ = 0.74 - 0.77, RMSE = 0.13 - 0.30 in training, R^2^ = 0.57 - 0.76, RMSE = 0.14 - 0.39 in testing). Our RF model in the vegetative growth phase allowed the best prediction for spike and plant NC, respectively, in the training and testing sets. Similar to the PLSR model in the vegetative growth phase, the prediction of NC by the RF model did not show differences between different organs in wheat or the whole plant (R^2^ = 0.91 - 0.94, RMSE = 0.07 - 0.26 in training, R^2^ = 0.73 - 0.82, RMSE = 0.13 - 0.50 in testing). Figure 5 shows the PLSR and RF models that had the best overall performance in the vegetative and reproductive growth phases.

**Table 2.**
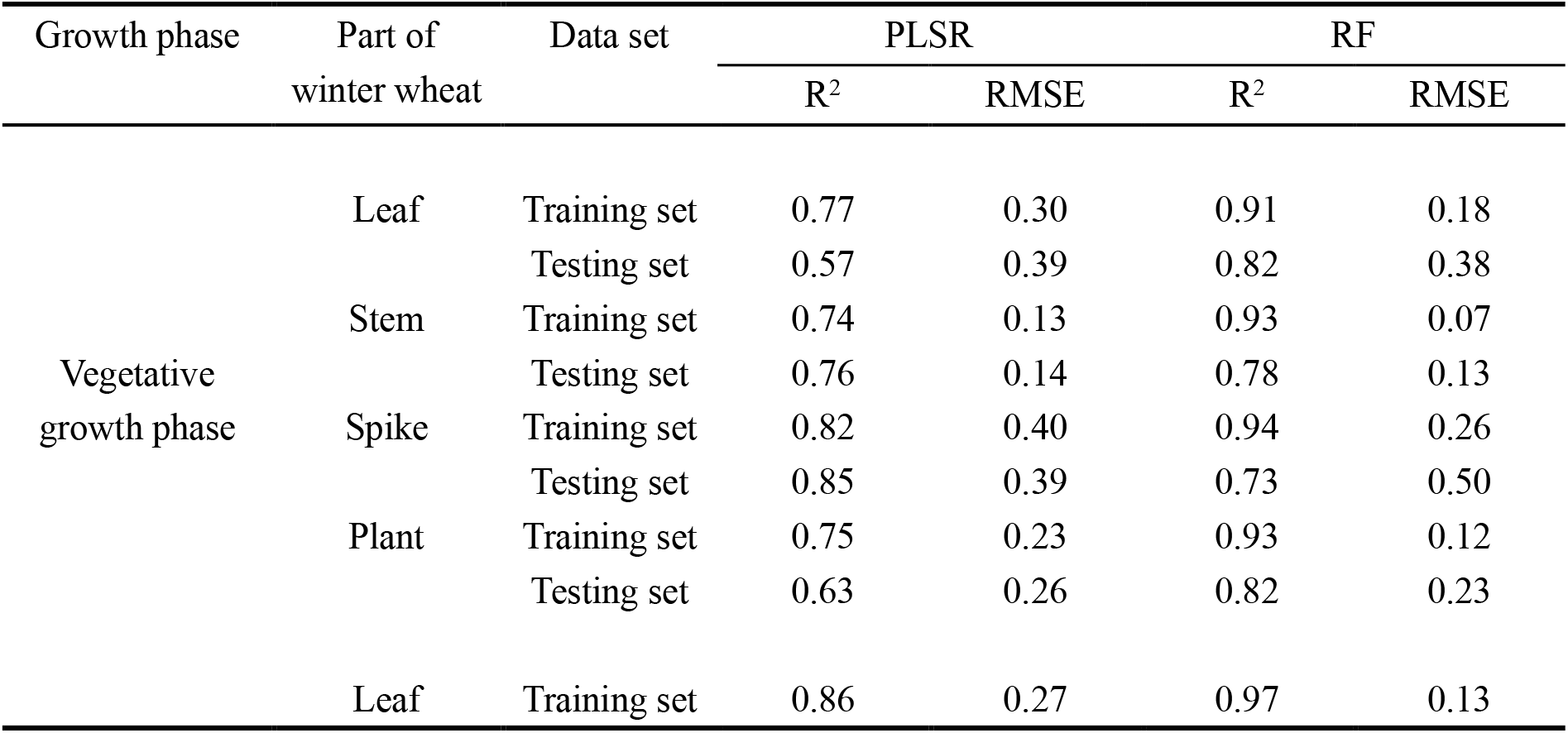
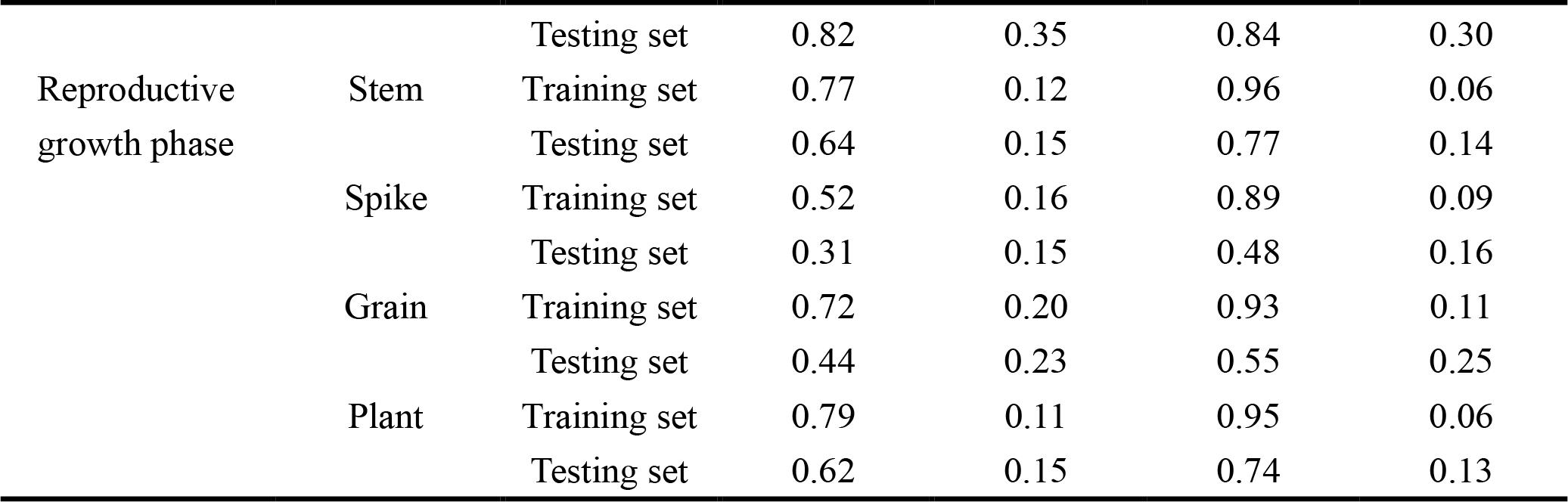
Nitrogen content estimates using 43 vegetation indices.

**Figure 5.**
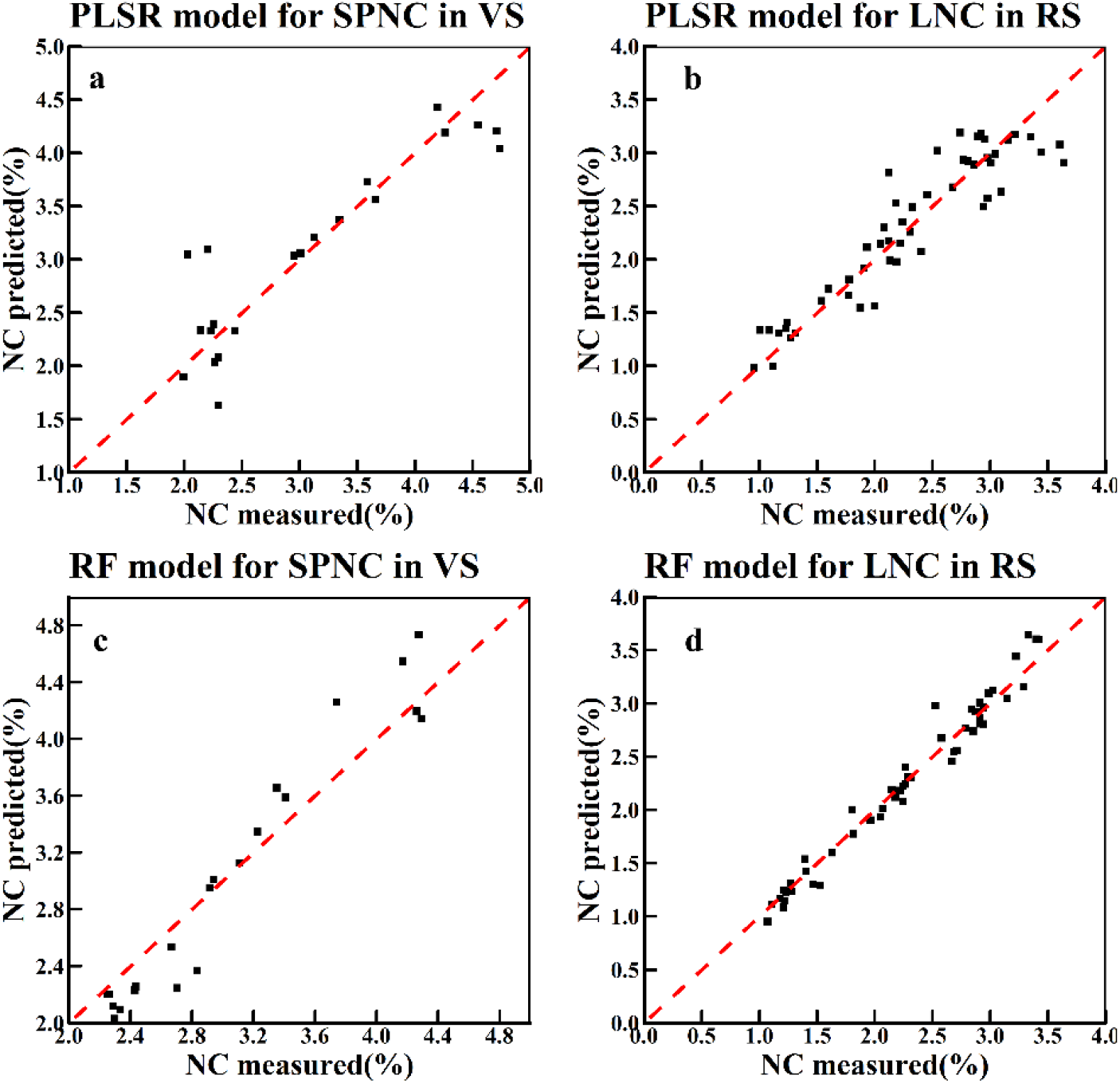
The PLSR and RF models which performed best in vegetative and reproductive growth phases using VIs only. (**a**) the SPNC PLSR model in VS. (**b**) the LNC PLSR model in RS. (**c**) the SPNC RF model in VS. (**d**) the LNC RF model in RS.

Figure 6 showed the top 10 important VIs for NC estimation models. Among all the NC of different organs or the whole plant in the vegetative growth phase, MCARI2 was found to be the most important VI for leaf NC (VIP = 1.98), spike NC (VIP = 4.87) and plant NC (VIP = 1.92) in PLSR models. MTCI was the 2nd most important VI for leaf NC (VIP = 1.51) and plant NC (VIP = 1.47) and was also found to be the most important VI for stem NC (VIP = 1.18).

**Figure 6.**
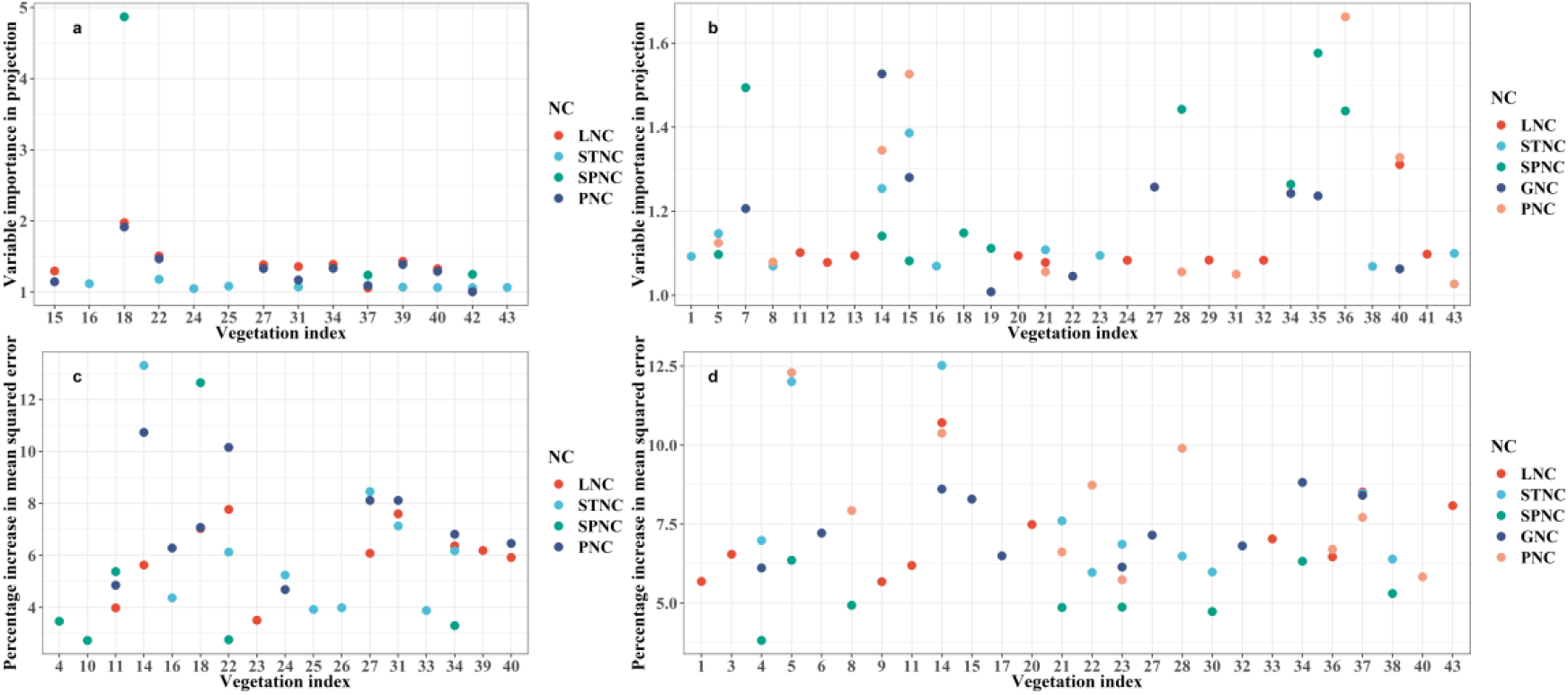
Top 10 important VIs for the NC monitoring of different organs and the whole plant selected by different models. (**a**) the TOP 10 important VIs for NC monitoring in vegetative growth phase selected by PLSR. (**b**) the TOP 10 important VIs for NC monitoring in the reproductive growth phase selected by PLSR. (**c**) the TOP 10 important VIs for NC monitoring in the vegetative growth phase selected by RF. (**d**) the TOP 10 important VIs for NC monitoring in the reproductive growth phase selected by RF. LNC, STNC, SPNC, GNC and PNC are leaf, stem, spike, grain and plant NC, respectively.

As for the RF models in the vegetative growth phase, MTCI, GRVI, MCARI2 and GRVI with contributed most to the leaf-, stem-, spike- and plant NC estimations, respectively, with the %IncMSE of 10.73, 13.31, 12.64 and 10.73 (Figure 6). Also, MCARI2 and MTCI also played an important role in the RF models, which had similarly great performance in the PLSR models during the vegetative growth.

For the reproductive growth phase, the VIs that yielded great performance in the NC prediction models differed. In the PLSR models. TCARI/OSAVI, LCI, SAVI-GREEN, GRVI and S-CCCI have been found to be the best VIs for leaf, stem, spike, grain and plant NC, respectively, with the VIP of 1.31, 1.37, 1.58, 1.52 and 1.66. In the RF models, GRVI contributed most to the leaf- and stem NC predictions (%IncMSE = 10.71 and 12.52), CVI contributed most to spike and plant NC (%IncMSE of 6.36 and 12.30), and SAVI contributed most for grain NC (%IncMSE = 8.82). Besides, it also indicated that the VIs screened out in the vegetative growth phase are more consistent, while weak consistency of the top 10 VIs in the reproductive growth phase (Figure 6). Furthermore, we have also counted the total number of VIs selected by the PLSR and RF models in different growth phases. Table S4 shows that more VIs have been selected by RF models in the reproductive growth phase of winter wheat approximately (See detail in Supplementary Material S3).

### 3.4 PLSR and RF models using texture features for nitrogen content estimation

In Table 3, it can be found that during the vegetative growth phase, both PLSR (R^2^ = 0.84, RMSE = 0.16) and RF (R^2^ = 0.97, RMSE = 0.06) model performed the best for plant NC estimation in the training set. And for leaf and spike NC estimation, both PLSR and RF models achieved great performance with R^2^ above 0.79, RMSE below 0.28 in the training set and R^2^ above 0.53, RMSE below 0.35 in the testing set. Besides, the results also showed that it was more stable for the prediction of leaf NC than stem NC since the worse performance of both PLSR and RF models in the testing set.

**Table 3.** Nitrogen content estimates using 40 texture features.

| Growth stage | Part of winter wheat | Data set | PLSR |  | RF |  |
| --- | --- | --- | --- | --- | --- | --- |
|  |  |  | R<sup>2</sup> | RMSE | R<sup>2</sup> | RMSE |
| Vegetative growth stage | Leaf | Training set | 0.79 | 0.28 | 0.94 | 0.14 |
|  |  | Testing set | 0.72 | 0.31 | 0.87 | 0.35 |
|  | Stem | Training set | 0.81 | 0.11 | 0.96 | 0.06 |
|  |  | Testing set | 0.53 | 0.21 | 0.77 | 0.13 |
|  | Spike | Training set | 0.57 | 0.34 | 0.97 | 0.18 |
|  |  | Testing set | 0.60 | 0.44 | 0.94 | 0.23 |
|  | Plant | Training set | 0.87 | 0.16 | 0.97 | 0.08 |
|  |  | Testing set | 0.72 | 0.24 | 0.90 | 0.17 |
| Reproductive growth stage | Leaf | Training set | 0.88 | 0.25 | 0.97 | 0.14 |
|  |  | Testing set | 0.88 | 0.32 | 0.76 | 0.38 |
|  | Stem | Training set | 0.73 | 0.13 | 0.94 | 0.07 |
|  |  | Testing set | 0.76 | 0.12 | 0.84 | 0.13 |
|  | Spike | Training set | 0.57 | 0.16 | 0.91 | 0.09 |
|  |  | Testing set | 0.16 | 0.18 | 0.31 | 0.19 |
|  | Grain | Training set | 0.66 | 0.23 | 0.93 | 0.12 |
|  |  | Testing set | 0.26 | 0.28 | 0.40 | 0.27 |
|  | Plant | Training set | 0.74 | 0.13 | 0.94 | 0.06 |
|  |  | Testing set | 0.68 | 0.14 | 0.74 | 0.13 |

In the reproductive growth phase, the performance of the PLSR (R^2^ = 0.88, RMSE = 0.25 in training, R^2^ = 0.88, RMSE = 0.32 in testing) and RF (R^2^ = 0.97, RMSE = 0.14 in training, R^2^ = 0.76, RMSE = 0.38 in testing) models for the leaf NC prediction were improved. However, the performance of the PLSR (R^2^ = 0.57, RMSE = 0.16 in training and R^2^ = 0.16, RMSE = 0.18 in testing) and RF (R^2^ = 0.91, RMSE = 0.09 in training, R^2^ = 0.31, RMSE = 0.19 in testing) models for the spike NC prediction was worse than that in the vegetative growth phase. For plant and stem NC monitoring, no significant differences were found between two different stages. Besides, the prediction of grain NC has achieved fairly good performance in the training set though it did not allow great performance in the testing set. We can find the PLSR and RF models with the best overall performance in the vegetative and reproductive growth phases in figure 7.

**Figure 7.**
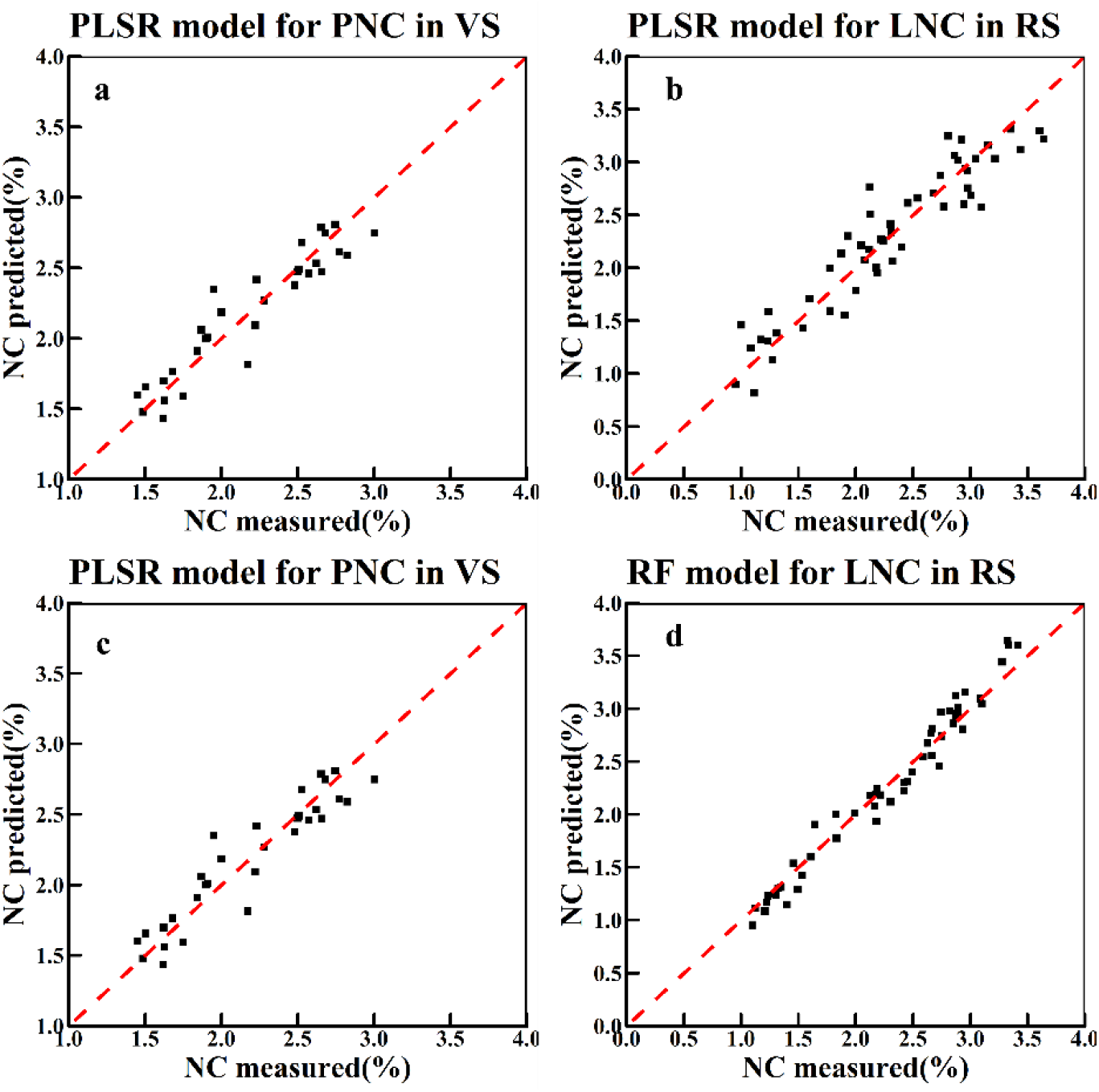
The PLSR and RF models which performed best in vegetative and reproductive growth phases using TFs only. (**a**) the PNC PLSR model in VS. (**b**) the LNC PLSR model in RS. (**c**) the PNC RF model in VS. (**d**) the LNC RF model in RS.

Figure 8 shows the top 10 important TFs for NC estimation models except for the RF models for spike and plant NC for there were fewer than 10 TFs were screened. In the vegetative growth phase, The best TF for leaf, stem, spike and plant NC were Reg_mean (VIP = 1.41), G_cor (VIP = 1.43), B_cor (VIP = 1.29) and Reg_mean (VIP = 1.42) for the PLSR models. For RF models, B_mean with %IncMSE of 10.43, R_mean with %IncMSE of 12.69, G_mean with %IncMSE of 7.66 and G_mean with %IncMSE of 11.90 was the best TFs for the estimating of leaf, stem, spike and plant NC, respectively. In the reproductive growth phase, for PLSR models, R_ho, G_mean, Reg_cor, B_cor and Reg_mean have been found to be the best TFs for leaf, stem, spike, grain and plant NC, respectively, with the VIP of 1.32, 1.41, 1.40, 1.40 and 1.47. In contrast in the RF models, R_dis, G_mean, G_con and contributed the most to the leaf-, stem-, and grain NC preditions, respectively, with the %IncMSE of 12.04, 8.38 and 7.75. B_cor performed the best for spike and plant NC predictions, respectively, with %IncMSE of 10.52 and 13.22. Furthermore, the result also indicated that the TFs of mean and cor accounted for a relatively large proportion of the variations in both PLSR and RF models.

**Figure 8.**
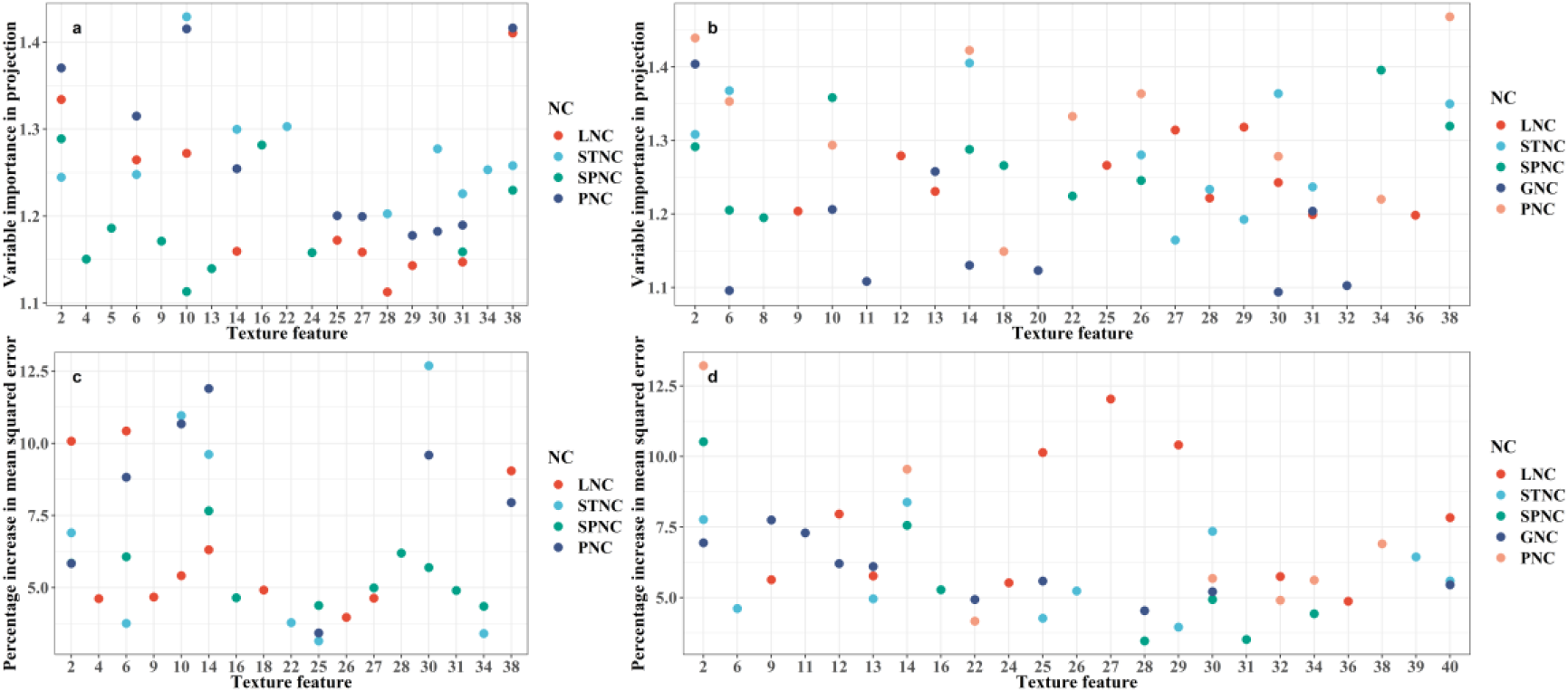
Top 10 important TFs for the NC monitoring of different organ and the whole plant selected by different models. (**a**) the TOP 10 important TFs for NC monitoring in vegetative growth phase selected by PLSR. (**b**) the TOP 10 important TFs for NC monitoring in reproductive growth phase selected by PLSR. (**c**) the TOP 10 important TFs for NC monitoring in vegetative growth phase selected by RF. (**d**) the TOP 10 important TFs for NC monitoring in reproductive growth phase selected by RF. LNC, STNC, SPNC, GNC and PNC are leaf, stem, spike, grain and plant NC, respectively.

Table S5 shows the number of TFs selected by the PLSR and RF models in different growth phases. It can be found that more TFs were selected by the PLSR models than the RF models. Meanwhile, by counting the TFs screened by the two models, it was found that almost all the important TFs screened out by the models were based on the bands of R, G and B instead of NIR and REG bands (See detail in Supplementary Material S2).

### 3.5 PLSR and RF models using the combination of VIs and texture features for nitrogen content estimation

Table 4 showed that the combination of image VIs and TFs did improve the monitoring accuracy of NC in winter wheat to a certain extent, but the effect was not significant. Among all the models in the vegetative growth phase, the estimation for spike NC has allowed great performance in both PLSR (R^2^ = 0.93, RMSE = 0.25 in training and R^2^ = 0.77, RMSE = 0.33 in testing) and RF (R^2^ = 0.98, RMSE = 0.16 in training and R^2^ = 0.94, RMSE = 0.23 in testing) models. And better results have been achieved for plant NC monitoring than leaf stem NC monitoring (R^2^ = 0.82 - 0.87, RMSE = 0.06 - 0.26 in the training set, R^2^ = 0.52 - 0.94, RMSE = 0.17 - 0.40 in testing set).

**Table 4:**
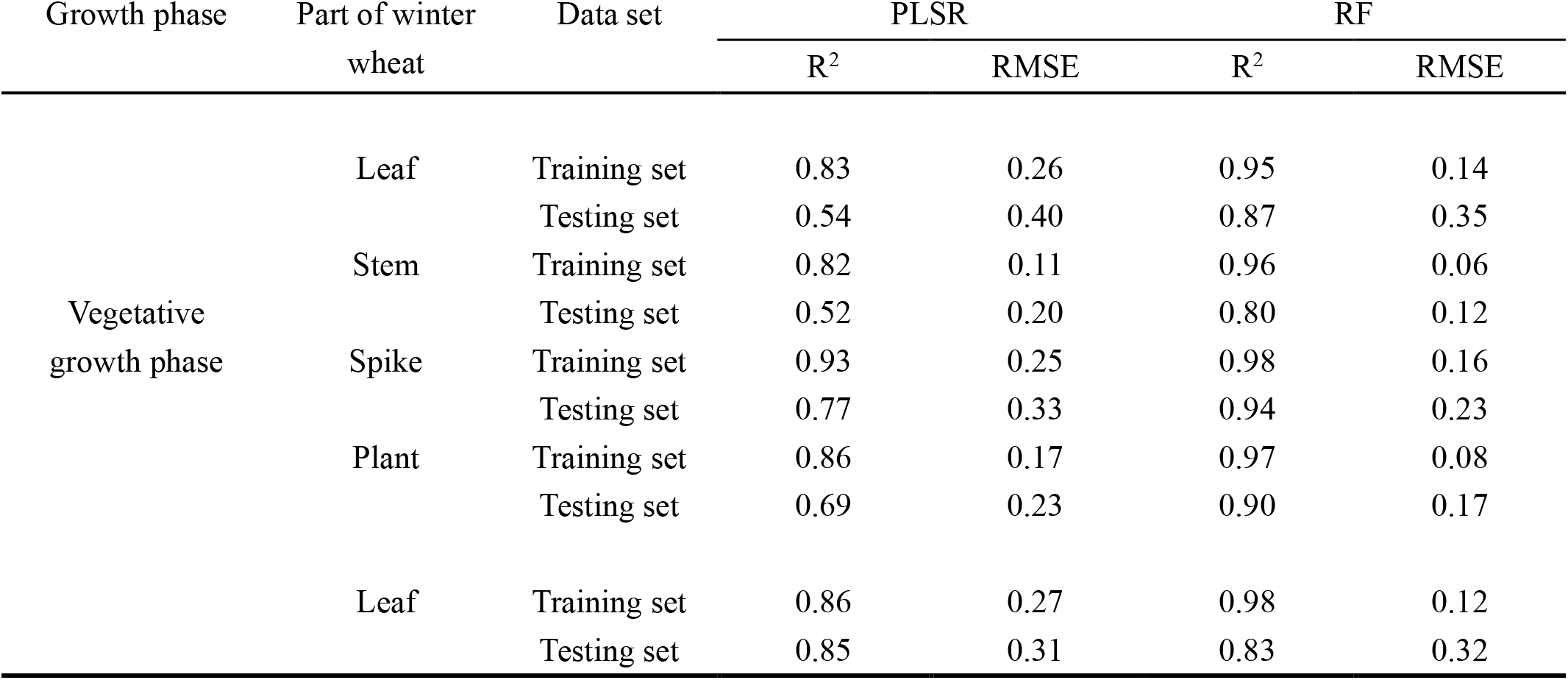
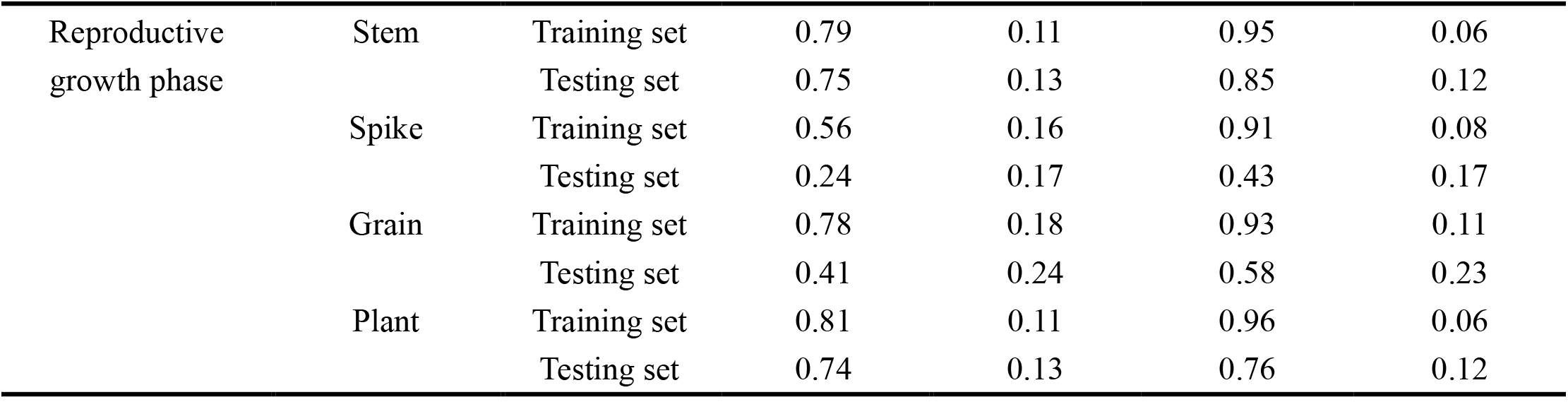
Nitrogen content estimates using the combination of vegetation indices and texture features.

| Growth phase | Part of winter wheat | Data set | PLSR |  | RF |  |
| --- | --- | --- | --- | --- | --- | --- |
|  |  |  | R<sup>2</sup> | RMSE | R<sup>2</sup> | RMSE |
| Vegetative growth phase | Leaf | Training set | 0.83 | 0.26 | 0.95 | 0.14 |
|  |  | Testing set | 0.54 | 0.40 | 0.87 | 0.35 |
|  | Stem | Training set | 0.82 | 0.11 | 0.96 | 0.06 |
|  |  | Testing set | 0.52 | 0.20 | 0.80 | 0.12 |
|  | Spike | Training set | 0.93 | 0.25 | 0.98 | 0.16 |
|  |  | Testing set | 0.77 | 0.33 | 0.94 | 0.23 |
|  | Plant | Training set | 0.86 | 0.17 | 0.97 | 0.08 |
|  |  | Testing set | 0.69 | 0.23 | 0.90 | 0.17 |
|  | Leaf | Training set | 0.86 | 0.27 | 0.98 | 0.12 |
|  |  | Testing set | 0.85 | 0.31 | 0.83 | 0.32 |
| Reproductive<br>growth phase | Stem | Training set | 0.79 | 0.11 | 0.95 | 0.06 |
|  |  | Testing set | 0.75 | 0.13 | 0.85 | 0.12 |
|  | Spike | Training set | 0.56 | 0.16 | 0.91 | 0.08 |
|  |  | Testing set | 0.24 | 0.17 | 0.43 | 0.17 |
|  | Grain | Training set | 0.78 | 0.18 | 0.93 | 0.11 |
|  |  | Testing set | 0.41 | 0.24 | 0.58 | 0.23 |
|  | Plant | Training set | 0.81 | 0.11 | 0.96 | 0.06 |
|  |  | Testing set | 0.74 | 0.13 | 0.76 | 0.12 |

In the reproductive growth phase, the worst performance was obtained when estimating spike NC (R^2^ = 0.56, RMSE = 0.16 in the training set and R^2^ = 0.24, RMSE = 0.17 in the testing set for PLSR model and R^2^ = 0.91, RMSE = 0.08 in the training set and R^2^ = 0.43, RMSE = 0.17 in the testing set for RF model). The performance of grain NC was also not so satisfactory in testing set, with R^2^ of 0.41 and RMSE of 0.24 in the PLSR model and R^2^ of 0.43 and RMSE of 0.17 in the RF model. Apart from that, the best performance was achieved in leaf NC prediction with the highest R^2^ of 0.86 in PLSR model and 0.98 in RF model. Figure 9 shows the PLSR and RF models with the best overall performance in the vegetative and reproductive growth phases.

**Figure 9.**
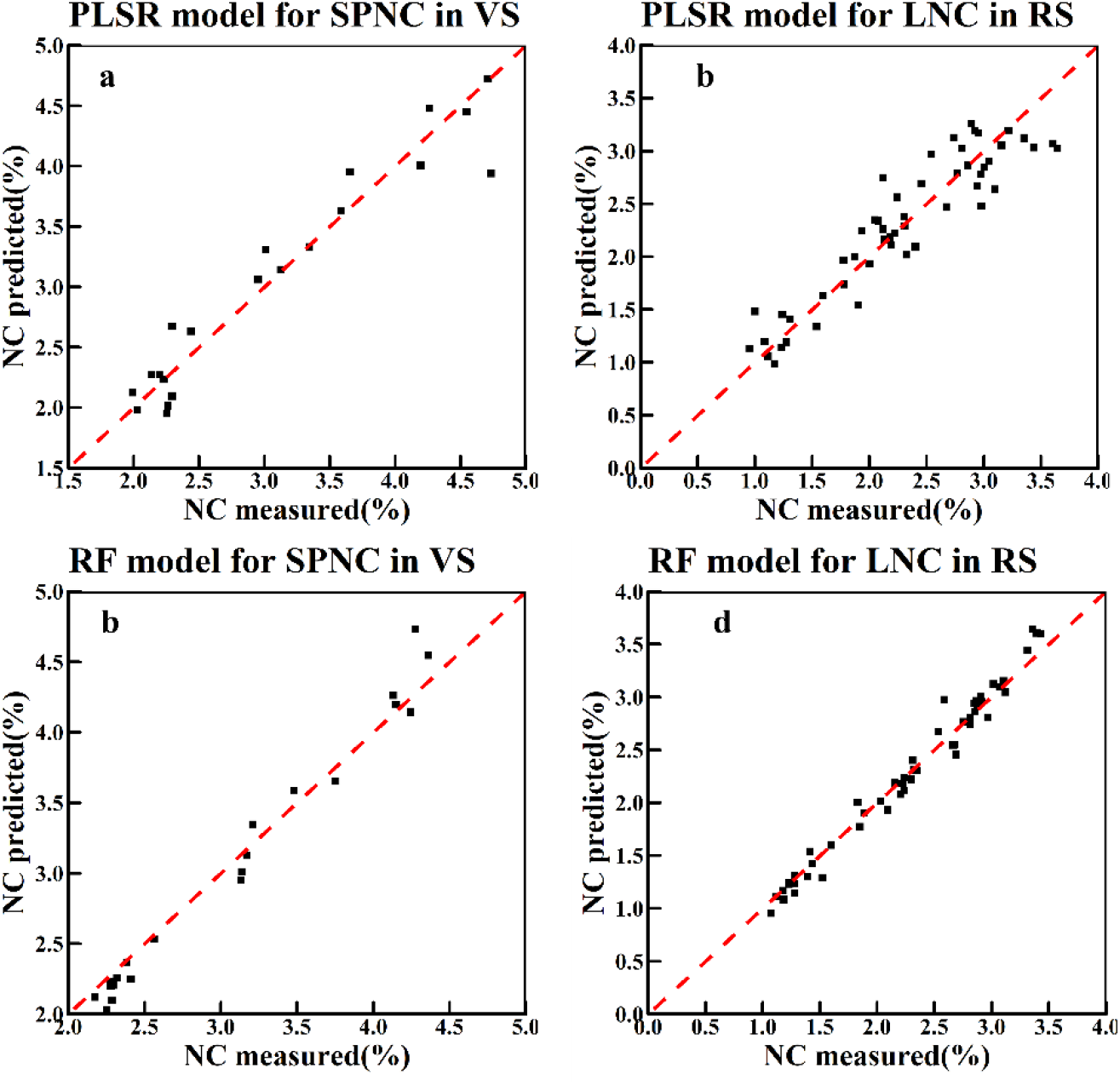
The PLSR and RF models which performed best in vegetative and reproductive growth phases using TFs only. (**a**) the SPNC PLSR model in VS. (**b**) the LNC PLSR model in RS. (**c**) the SPNC RF model in VS. (**d**) the LNC RF model in RS.

Figure 10 shows the TOP 10 selected features based on the PLSR and RF models. In the vegetative growth, the feature with the highest VIP was Reg_mean (VIP = 1.51) for leaf NC, G_cor (VIP = 1.32) for stem NC, MCARI2 (VIP = 1.66) for spike NC and Reg_mean (VIP = 1.43) for palnt NC in PLSR model. In RF model, B_mean with %IncMSE of 9.26, R_mean with %IncMSE of 10.54, G_mean with %IncMSE of 7.53 and 12.39 was the best feature for leaf, stem, spike and plant NC. In the reproductive growth stage, GOSAVI, Reg_mean, SAVI-GREEN, GRVI and Reg_mean (with VIP of 1.24, 1.36, 1.39, 1.60 and 1.45) contributed most to the leaf, stem, spike, grain and plant NC in PLSR models. R_var, GRVI, B_cor, SAVI and B_cor (with %IncMSE of 9.79, 7.94, 8.01, 7.72 and 11.29) contributed most to the corresponding NC estimation in RF models. Compared with the best image features selected in different growth phases of winter wheat, it also reflected that the TFs could be more suitable for the monitoring of NC of winter wheat in general. As for the total number of image features (VIs and TFs) selected by the PLSR and RF models in different growth phases. For all the PLSR and RF models except for STNC, more TFs was screened than VIs in the vegetative growth phase. Interestingly, in the reproductive growth phase, more VIs were screened out than TFs in all the PLSR and RF models except for SPNC, which was different from the characteristics of in the vegetative growth phase (See detail in Supplementary Material S2).

**Figure 10.**
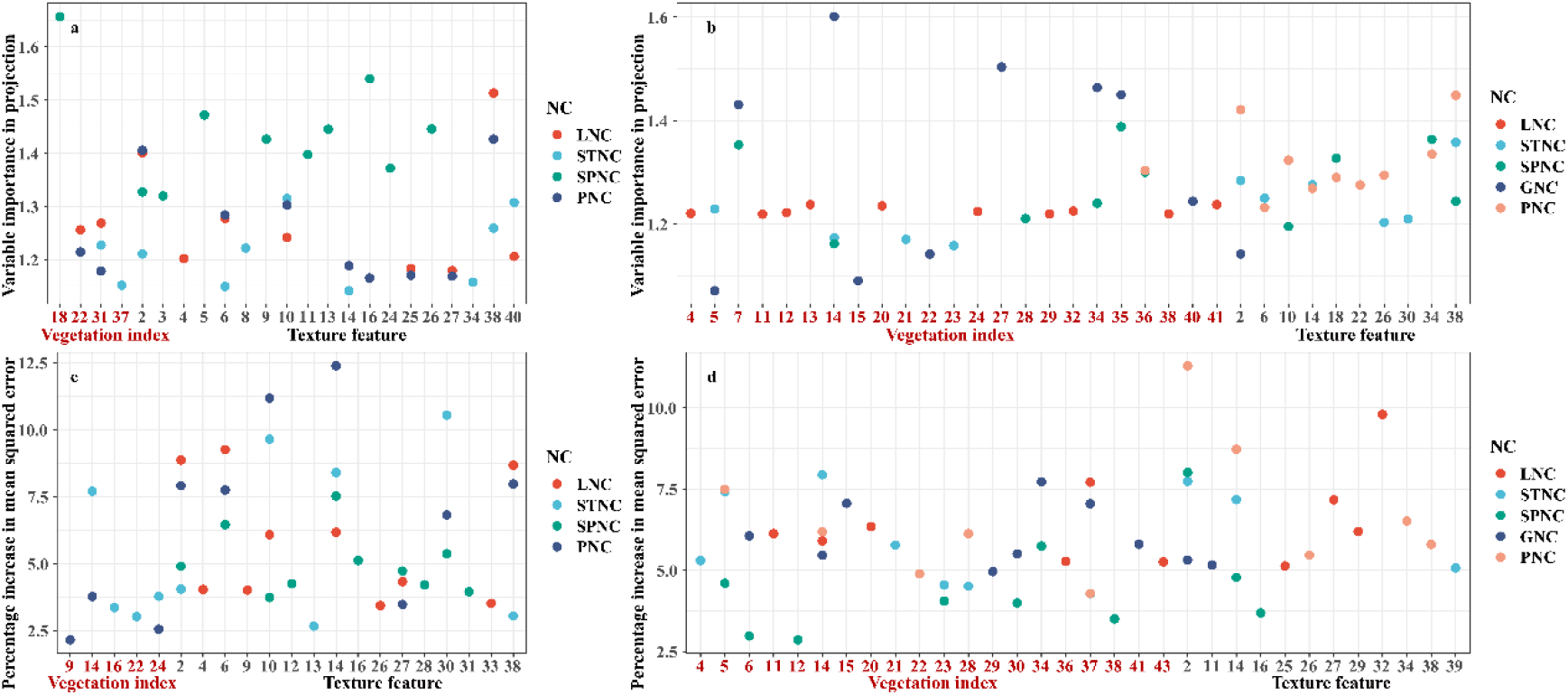
Top 10 important image features (VIs and TFs) for the NC monitoring of different organs and the whole plant selected by different models. (**a**) the TOP 10 important image features for NC monitoring in the vegetative growth phase selected by PLSR. (**b**) the TOP 10 important image features for NC monitoring in the reproductive growth phase selected by PLSR. (**c**) the TOP 10 important image features for NC monitoring in vegetative growth phase selected by RF. (**d**) the TOP 10 important image features for NC monitoring in the reproductive growth phase selected by RF. LNC, STNC, SPNC, GNC and PNC are leaf, stem, spike, grain and plant NC, respectively.

## 4 Discussion

### 4.1 UAV-based predictions of nitrogen content in organs and whole plants of winter wheat

In this study, during vegetative and reproductive growth phases, not only the correlation between image features (VIs and TFs) and NC of winter wheat were analyzed, but also the corresponding PLSR and RF models were constructed for the different organs or the whole plant of winter wheat. As found in several preview studies (Zheng et al., 2018; Fu et al., 2020), the leaf and plant NC can be well estimated using VIs or TFs derived from UAV-based images. Our study has also shown great performance of both types of variables for leaf and plant NC predictions during the vegetative and reproductive growth phases. It is worth noting that these variables extracted from the images obtained from the UAV have the capability of estimating the stem, spike and grain NC.

It is worth noting that the spike NC always yielded the lowest correlations with VIs and TFs when compared to other organs or the whole plant (Table 3) and, that the predictions for spike NC were not as satisfactory as that for other organs. In contrast, the leaf-, stem- and plant NC were highly correlated in different growth stages, especially in the reproductive growth phase (Figure 11). The relatively low correlations in the vegetative growth phase suggest that the rapid changes in canopy structure during the vegetative growth phase constrained the predictions for leaf, stem and plant NC (Yu et al., 2014). In this study, the VIs and TFs were derived from the delineated subplots (about 30 m^2^), which reflected the spectral reflectance as a response to the crop canopy variations. Compared to spikes, it is certain that, in orthophotos acquired by the UAV, leaves contributed relatively large to the canopy spectrum (Liu et al., 2017; Yang et al., 2021), which may explain the relatively weak correlations with the extracted VIs and TFs and the relatively high predictions errors (RMSE) for spike NC.

**Figure 11.**
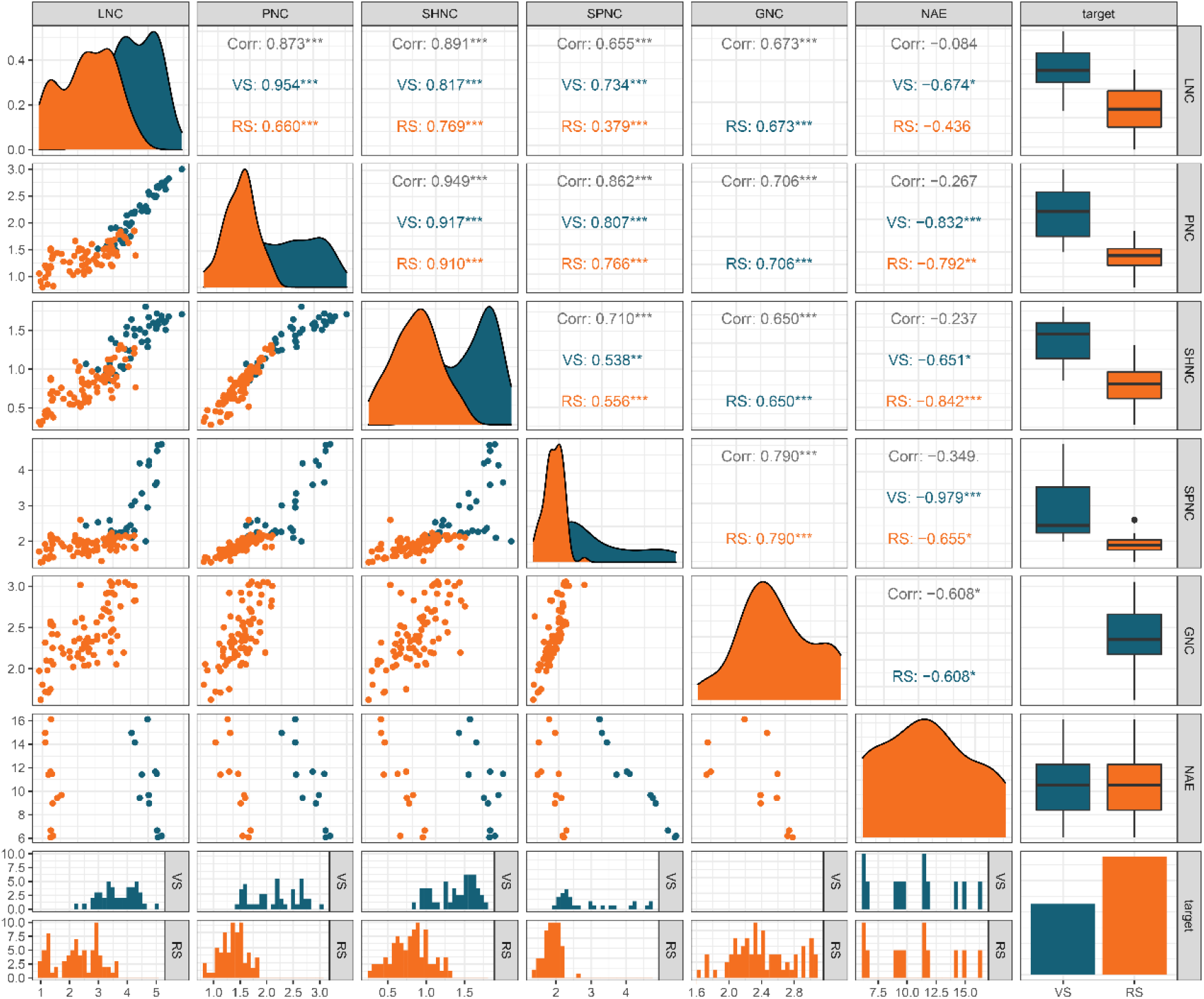
Correlation between nitrogen content and NAE from different organs or the whole plant of winter wheat. LNC, STNC, SPNC, GNC and PNC are leaf, stem, spike, grain and plant NC, respectively. NAE is the nitrogen agronomic efficiency. VS and RS means vegetative and reproductive growth phases. NAE are correlated with the NC of different organs or the whole plant obtained from two stages (booting and heading stage) in VS, and five stages (AF5, AF10, AF15, AF20, AF25) in RS.

### 4.2 Comparisons between the vegetative and reproductive growth phases

Many studies have raised the importance of growth stage on crop agronomic parameters monitoring (Xue et al., 2004; Li et al., 2010; Wang et al., 2019) found the leaf and plant NC could be well predicted during the vegetative growth phase including tillering, jointing, booting and heading stages of rice. Similar studies revealed the monitoring performance of leaf NC for winter wheat in the reproductive growth phase could be worse than it is performed in vegetative growth phase (Zheng et al., 2018; Ge et al., 2021; Wang et al., 2022b).

In contrast, our results showed inconsistency regarding the best growth stages for leaf NC prediction. Based on our PLSR and RF models, better prediction performance could be achieved for predicting leaf NC in the reproductive growth phase though predicting leaf NC in the vegetative growth phase was also successful. This is attributed to the fact that the unclosed canopy and soil would be the confusing factors for canopy reflectance in the early vegetative growth phase (Li et al., 2010). Also, the large variations in biomass over early growth stages will also be responsible for the worse performance of leaf NC prediction (Yu et al., 2013). In addition, the prediction of spike NC was found to have the opposite trend compared to the leaf NC, i.e., the vegetative growth phase allowed the best prediction of spike NC. As the reproductive organ of winter wheat, the spike acts as a major photosynthetic organ during the grain filling and has great relevance for plant nitrogen assimilation (Sanchez-Bragado et al., 2014; Vicente et al., 2018). Recent studies have revealed that spikes have certain effects on canopy reflectance spectra, though the complexity of canopy structure, plant density and morphoanatomical and compositional characteristics of spikes in response to canopy spectra still needs to be investigated (Li et al., 2015; Vergara-Diaz et al., 2020).

After reaching the reproductive growth phase, the grain appears and becomes the “growth center” of the plant; the N transport mainly happens from the leaf, stem, glume and awn to grain (Maydup et al., 2012; Sanchez-Bragado et al., 2016; Vergara-Diaz et al., 2020). The bad performance of grain NC using PLSR and RF models indicated that grain could be the major confusing factor for the bad performance of spike NC monitoring in the reproductive growth phase, since we could not fully capture the spectral information of grain which was wrapped in glume. Furthermore, compared with leaf, the delayed senescence of spike may also worsen the performance for spike NC monitoring in the reproductive growth phase (Kong et al., 2015; Vicente et al., 2018). However, no significant differences have been found between the two growth phases for the plant ant stem NC predictions, which does not allow us to conclude on which stages could be more suitable for the whole plant and stem NC estimation.

### 4.3 Comparison between image feature types (VIs and TFs)

Our result has shown that both VIs and TFs can be great features for winter wheat N monitoring. However, inconsistent with the results which were highlighted in crop biomass monitoring (Yue et al., 2019; Zheng et al., 2019), the combination of VIs and TFs didn’t significantly improve the estimation accuracy of NC of winter wheat in our study. Actually, there were a few studies focused on the contribution of the integration of VIs and TFs for crop N monitoring and generally, they concluded that combining VIs and TFs performed better than only using the VIs or TFs, e.g., for leaf and plant NC monitoring (Jia and Chen, 2020; Zheng et al., 2020). The multiple types of VIs can make more extensive use of waveband information and provide more complementary predictors for the NC model construction. Thus, the machine learning algorithms have the ability to integrate and utilize the spectral information contained in VIs, which could be the explanation for the great performance achieved for the combined use of VIs (Wang et al., 2022a). However, probably due to the contrasting correlation patterns observed here - VIs and TF were correlated positively and negatively with NC respectively, the combined use of both types of variables did not improve the predictions of NC.

By comparing screened image features, there are a few interesting patterns that deserve our attention. Firstly, compared to the image features screened out in the vegetative growth phase (Figures 6, 8, 10), more features with strong consistency were screened out for the PLSR and RF models of different organs in the reproductive growth phase. This could be explained by the complicated canopy structure of winter wheat in the late growth stages, leading to many problems for crop monitoring, such as the saturated VIs (Haboudane et al., 2004). Secondly, among all the top 10 VIs screened out for different organs, most VIs such as MCARI2, MTCI, TCARI, TCARI/OSAVI, SAVI and OSAVI could fall into the ‘soil-line’ VIs and the VIs related to chlorophyll. For example, MCARI2 was reported to be the sensitive VI for the monitoring of N status in the early stage of maize and winter wheat (Nigon et al., 2020). MTCI have also been reported to be the promising spectral index for determining N stress level of potato (Nigon et al., 2015), monitoring the leaf NC of rice (Tian et al., 2011) and estimating the N status of maize (Li et al., 2014). As for the soil-line VIs, lots of studies have demonstrated its’ promise for N monitoring (Gabriel et al., 2017; Klem et al., 2018; Guo et al., 2019). The high correlation between N and chlorophyll and the strong ability to minimize soil background influence may be the main reason for the great performance of these indices in the early growth stages. In contrast, the VIs selected in the reproductive growth phase were not as consistent as they were in the vegetative growth phase. Thirdly, the result of selected TFs showed that among all the TFs derived from five different band, more TFs based on R, G and B band were selected by our PLSR and RF models. Also, the texture *mean* and *cor* features accounted for a large proportion in the selected top 10 TFs. It has been know that the mean and cor exhibited great performance in classification tasks (Wan and Chang, 2019). Similar results have been reported for the performance of the texture mean for biomass monitoring in (Fu et al., 2021). The texture mean reflects the degree of regularity of the texture and cor describes the similarity of elements within a line or a row in the GLCM features (Zhu et al., 2022), and thus it has the capability of smoothing the image and minimizing the interference of background. Lastly, although the performance of the combination of VIs and TFs did not show better performance for N monitoring compared with the models based only on VIs and TFs, the top 10 image features filtered by our models based on the combination of VIs and TFs indicated that TFs deserve more attention in the future research since more TFs were selected among the top 10 image features in almost all the models. Overall, these TFs should be further evaluated in future research, such as whether the accuracy of the models can be improved when using the normalized texture index or when monitoring nitrogen in different crop species and varieties.

### 4.4 UAV-based predictions of N use efficiency

As an important indication for crop N use efficiency, the potential of NAE for crop N status monitoring has not been well evaluated using UAV-based imaging. There were only limited studies reported the attempts on the UAV-based estimation of N use efficiency, which for instance is reflected by the correlation between the UAV-based multispectral traits with NUE (Yang et al., 2020). (Liang et al., 2021) has revealed the capability of using UAV multispectral imagery for the identification of high N use efficiency phenotype in rice. Our results demonstrated that, by only using the latent variables extracted from UAV images, we could predict the NAE (Figure 12), highlighting the prospect of using of UAV-based images to estimate the indicators of NUE. The results of Pearson’s correlation analysis (Figure 4) over growth stags also confirm the findings of previous studies that the VIs derived from the multi-temporal images have the potential to forecast the canopy growth dynamics in relation to NUE. Also, the relatively better correlations between NC and NAE in the vegetative growth phase (Figure 11) than in the reproductive growth phase suggest the potential of assessing NUE in the early stages, e.g., for crop variety testing purposes.

**Figure 12.**
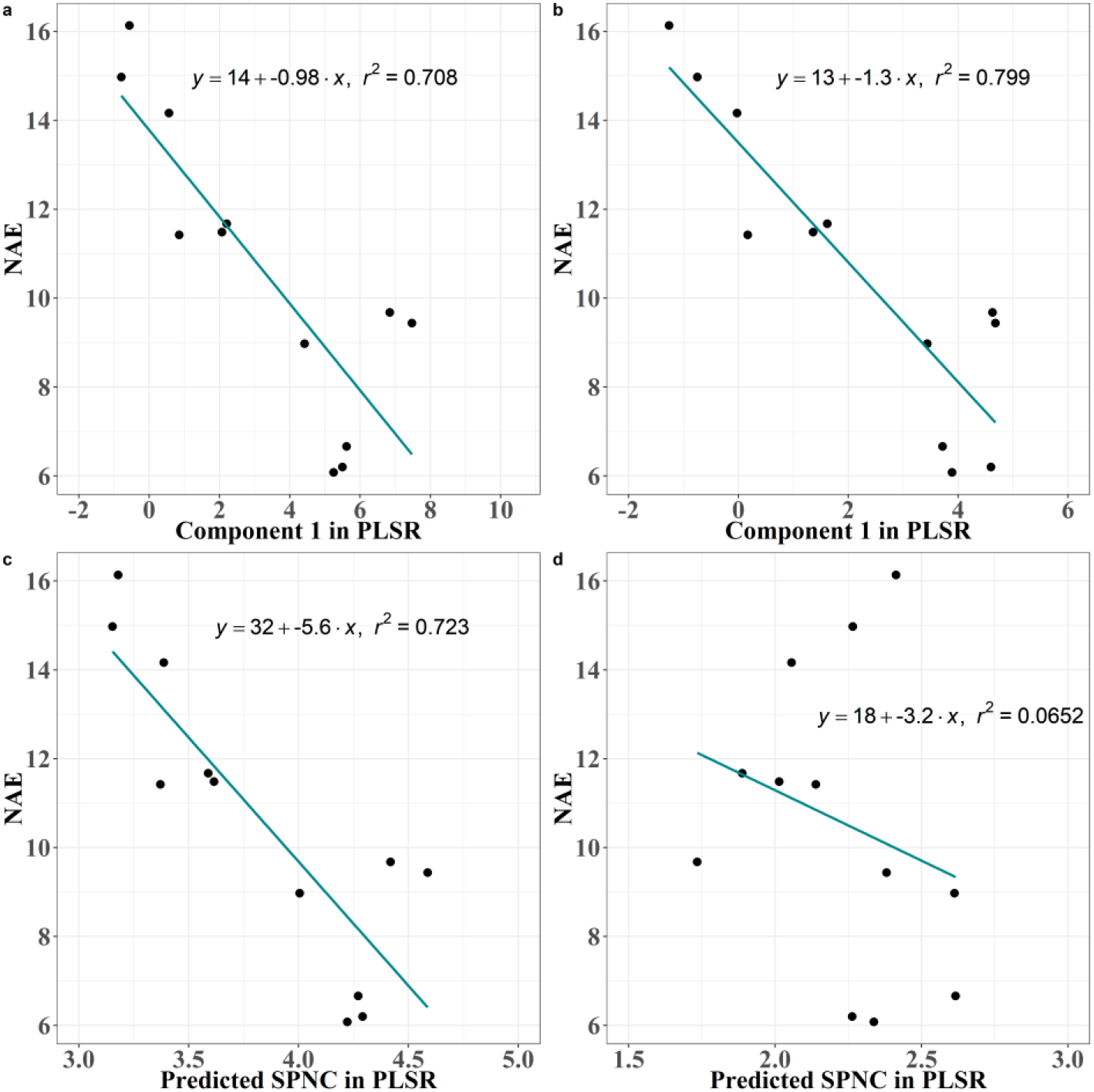
The performance of using the ‘Component 1’ and the predicted SPNC from the PLSR model in the vegetative growth phase for NAE predicting. (a) the performance of using the component 1 in the PLSR model for NAE predicting in the booting stage; (b) the performance of using the component 1 in the PLSR model for NAE predicting in the heading stage; (c) the performance of using the predicted SPNC in the PLSR model for NAE predicting in booting stage; (d) the performance of using the predicted SPNC in the PLSR model for NAE predicting in heading stage.

Furthermore, since the NAE is derived from the yield, the high correlation between VIs and NAE might also be due to the observed better performance for spike NC predictions in the vegetative growth phase. It is worth noting that the application of N fertilizer of winter wheat is mainly in the early growth stages during the vegetative growth phase, and thus the accurate monitoring of wheat N status in the early growth stage will provide more practical implications for wheat N fertilization for improved NUE and reduced environmental costs.

## 5 Conclusions

In this study, the muti-temporal measured nitrogen content (NC) in different organs or the whole plant of winter wheat obtained by field sampling was associated with the corresponding images acquired by a muti-spectral UAV. Stem-, spike- and plant-NC are found to decrease as dry matter weight (DMW) increased. Positive correlations were found between most of the VIs and NC, while negative correlations were found between most of the TFs and NC. PLSR and RF models successfully employed the VIs, TFs and their combinations to estimate the NC in the whole plant and different organs. PLSR latent variables extracted from the VIs and TFs explained successfully predicted the nitrogen agronomic efficiency (NAE). Although no significant differences were found between the VIs and TFs in their performance in predicting NC, some VIs like MCARI2 and TFs like texture mean were found to perform well in predicting NC. Finally, this study demonstrates that it is feasible to use UAV imaging and PLS/RF models to estimate NC and nitrogen use efficiency both in the vegetative and reproductive growth phases of winter wheat.

## DATA AVAILABILITY STATEMENT

The datasets generated for this study are available on request to the corresponding author.

## AUTHOR CONTRIBUTIONS

Experiments were designed by F.W. and K.Y.; F.W., Y.L., L.M., and L.Q performed the flight missions and completed the acquisition of dry matter weight of winter wheat in the field; F.W. compiled the data and conducted the data analysis; W.L. provided software technical support. Z.W., Y.Z., Z.S. and K.Y. supervised the experiments; F.W. wrote the initial draft of the manuscript and F.L. and K.Y. revised and edited the manuscript. All authors have approved the submitted version of the manuscript.

## ACKNOWLEDGMENTS

We thank the Wuqiao Experimental Station of China Agricultural University for the experiment site and equipment. We are also grateful for Ying Liu, Chenhang Du and Chunsheng Yao for their supports in field sampling. K.Y. appreciates the support by China Agricultural University while he was working at CAU.

## References

A., L. P. (1982). Methods of soil analysis. Part 2. Chemical and microbiological properties.

Bendig, J., Yu, K., Aasen, H., Bolten, A., Bennertz, S., Broscheit, J., et al. (2015). Combining UAV-based plant height from crop surface models, visible, and near infrared vegetation indices for biomass monitoring in barley. Int. J. Appl. Earth Obs. Geoinf. 39, 79–87. doi: 10.1016/j.jag.2015.02.012.

Berger, K., Verrelst, J., Féret, J. B., Wang, Z., Wocher, M., Strathmann, M., et al. (2020). Crop nitrogen monitoring: Recent progress and principal developments in the context of imaging spectroscopy missions. Remote Sens. Environ. 242, 111758. doi: 10.1016/j.rse.2020.111758.

Breiman, L., and Cutler, A. (2012). Breiman and Cutler’s random forests for classification and regression. Packag. “randomForest,” 29. Available at: https://cran.r-project.org/web/packages/randomForest/randomForest.pdf.

Cao, Q., Miao, Y., Wang, H., Huang, S., Cheng, S., Khosla, R., et al. (2013). Non-destructive estimation of rice plant nitrogen status with Crop Circle multispectral active canopy sensor. F. Crop. Res. 154, 133–144. doi: 10.1016/j.fcr.2013.08.005.

Chlingaryan, A., Sukkarieh, S., and Whelan, B. (2018). Machine learning approaches for crop yield prediction and nitrogen status estimation in precision agriculture: A review. Comput. Electron. Agric. 151, 61–69. doi: 10.1016/j.compag.2018.05.012.

Cui, Z., Zhang, F., Chen, X., Miao, Y., Li, J., Shi, L., et al. (2008). On-farm evaluation of an in-season nitrogen management strategy based on soil Nmin test. F. Crop. Res. 105, 48–55. doi: 10.1016/j.fcr.2007.07.008.

Errecart, P. M., Agnusdei, M. G., Lattanzi, F. A., and Marino, M. A. (2012). Leaf nitrogen concentration and chlorophyll meter readings as predictors of tall fescue nitrogen nutrition status. F. Crop. Res. 129, 46–58. doi: 10.1016/j.fcr.2012.01.008.

Farrés, M., Platikanov, S., Tsakovski, S., and Tauler, R. (2015). Comparison of the variable importance in projection (VIP) and of the selectivity ratio (SR) methods for variable selection and interpretation. J. Chemom. 29, 528–536. doi: 10.1002/cem.2736.

Fu, Y., Yang, G., Li, Z., Song, X., Li, Z., Xu, X., et al. (2020). Winter wheat nitrogen status estimation using uav-based rgb imagery and gaussian processes regression. Remote Sens. 12, 1–27. doi: 10.3390/rs12223778.

Fu, Y., Yang, G., Song, X., Li, Z., Xu, X., Feng, H., et al. (2021). Improved estimation of winter wheat aboveground biomass using multiscale textures extracted from UAV-based digital images and hyperspectral feature analysis. Remote Sens. 13, 1–22. doi: 10.3390/rs13040581.

Gabriel, J. L., Zarco-Tejada, P. J., López-Herrera, P. J., Pérez-Martín, E., Alonso-Ayuso, M., and Quemada, M. (2017). Airborne and ground level sensors for monitoring nitrogen status in a maize crop. Biosyst. Eng. 160, 124–133. doi: 10.1016/j.biosystemseng.2017.06.003.

Ge, H., Xiang, H., Ma, F., Li, Z., Qiu, Z., Tan, Z., et al. (2021). Estimating plant nitrogen concentration of rice through fusing vegetation indices and color moments derived from UAV-RGB images. Remote Sens. 13. doi: 10.3390/rs13091620.

Guo, C., Tang, Y., Lu, J., Zhu, Y., Cao, W., Cheng, T., et al. (2019). Predicting wheat productivity: Integrating time series of vegetation indices into crop modeling via sequential assimilation. Agric. For. Meteorol. 272–273, 69–80. doi: 10.1016/j.agrformet.2019.01.023.

Haboudane, D., Miller, J. R., Pattey, E., Zarco-Tejada, P. J., and Strachan, I. B. (2004). Hyperspectral vegetation indices and novel algorithms for predicting green LAI of crop canopies: Modeling and validation in the context of precision agriculture. Remote Sens. Environ. 90, 337–352. doi: 10.1016/j.rse.2003.12.013.

Hank, T. B., Berger, K., Bach, H., Clevers, J. G. P. W., Gitelson, A., Zarco-Tejada, P., et al. (2019). Spaceborne Imaging Spectroscopy for Sustainable Agriculture: Contributions and Challenges. Springer Netherlands doi: 10.1007/s10712-018-9492-0.

Haralick, R. M., Dinstein, I., and Shanmugam, K. (1973). Textural Features for Image Classification. IEEE Trans. Syst. Man Cybern. SMC-3, 610–621. doi: 10.1109/TSMC.1973.4309314.

Hastie, T., Tibshirani, R., and Friedman, J. (2005). The Elements of Statistical Learning : Data Mining, Inference and Prediction. Math. Intell. 27, 83–85. Available at: http://link.springer.com/article/10.1007/BF02985802?LI=true#.

Jia, D., and Chen, P. (2020). Effect of Low-altitude UAV Image Resolution on Inversion of Winter Wheat Nitrogen Concentration. Nongye Jixie Xuebao/Transactions Chinese Soc. Agric. Mach. 51, 164–169. doi: 10.6041/j.issn.1000-1298.2020.07.019.

Ju, X. T., Xing, G. X., Chen, X. P., Zhang, S. L., Zhang, L. J., Liu, X. J., et al. (2009). Reducing environmental risk by improving N management in intensive Chinese agricultural systems. Proc. Natl. Acad. Sci. U. S. A. 106, 3041–3046. doi: 10.1073/pnas.0813417106.

Kalacska, M., Lalonde, M., and Moore, T. R. (2015). Estimation of foliar chlorophyll and nitrogen content in an ombrotrophic bog from hyperspectral data: Scaling from leaf to image. Remote Sens. Environ. 169, 270–279. doi: 10.1016/j.rse.2015.08.012.

Kelsey, K. C., and Neff, J. C. (2014). Estimates of aboveground biomass from texture analysis of landsat imagery. Remote Sens. 6, 6407–6422. doi: 10.3390/rs6076407.

Kitonyo, O. M., Sadras, V. O., Zhou, Y., and Denton, M. D. (2018). Nitrogen supply and sink demand modulate the patterns of leaf senescence in maize. F. Crop. Res. 225, 92–103. doi: 10.1016/j.fcr.2018.05.015.

Klem, K., Záhora, J., Zemek, F., Trunda, P., Tůma, I., Novotná, K., et al. (2018). Interactive effects of water deficit and nitrogen nutrition on winter wheat. Remote sensing methods for their detection. Agric. Water Manag. 210, 171–184. doi: 10.1016/j.agwat.2018.08.004.

Kong, L., Sun, M., Xie, Y., Wang, F., and Zhao, Z. (2015). Photochemical and antioxidative responses of the glume and flag leaf to seasonal senescence in wheat. Front. Plant Sci. 6, 1–10. doi: 10.3389/fpls.2015.00358.

Lemaire, G., Jeuffroy, M. H., and Gastal, F. (2008). Diagnosis tool for plant and crop N status in vegetative stage. Theory and practices for crop N management. Eur. J. Agron. 28, 614–624. doi: 10.1016/j.eja.2008.01.005.

Li, D., Wang, X., Zheng, H., Zhou, K., Yao, X., Tian, Y., et al. (2018a). Estimation of area- and mass-based leaf nitrogen contents of wheat and rice crops from water-removed spectra using continuous wavelet analysis. Plant Methods 14, 1–20. doi: 10.1186/s13007-018-0344-1.

Li, F., Gnyp, M. L., Jia, L., Miao, Y., Yu, Z., Koppe, W., et al. (2008). Estimating N status of winter wheat using a handheld spectrometer in the North China Plain. F. Crop. Res. 106, 77–85. doi: 10.1016/j.fcr.2007.11.001.

Li, F., Miao, Y., Feng, G., Yuan, F., Yue, S., Gao, X., et al. (2014). Improving estimation of summer maize nitrogen status with red edge-based spectral vegetation indices. F. Crop. Res. 157, 111– 123. doi: 10.1016/j.fcr.2013.12.018.

Li, F., Miao, Y., Hennig, S. D., Gnyp, M. L., Chen, X., Jia, L., et al. (2010). Evaluating hyperspectral vegetation indices for estimating nitrogen concentration of winter wheat at different growth stages. Precis. Agric. 11, 335–357. doi: 10.1007/s11119-010-9165-6.

Li, H., Zhao, C., Yang, G., and Feng, H. (2015). Variations in crop variables within wheat canopies and responses of canopy spectral characteristics and derived vegetation indices to different vertical leaf layers and spikes. Remote Sens. Environ. 169, 358–374. doi: 10.1016/j.rse.2015.08.021.

Li, J., Shi, Y., Veeranampalayam-Sivakumar, A. N., and Schachtman, D. P. (2018b). Elucidating sorghum biomass, nitrogen and chlorophyll contents with spectral and morphological traits derived from unmanned aircraft system. Front. Plant Sci. 9, 1–12. doi: 10.3389/fpls.2018.01406.

Li, S., Ding, X., Kuang, Q., Ata-UI-Karim, S. T., Cheng, T., Liu, X., et al. (2018c). Potential of UAV-based active sensing for monitoring rice leaf nitrogen status. Front. Plant Sci. 871, 1–14. doi: 10.3389/fpls.2018.01834.

Li, W., Zhou, X., Yu, K., Zhang, Z., Liu, Y., Hu, N., et al. (2021). Spectroscopic Estimation of N Concentration in Wheat Organs for Assessing N Remobilization Under Different Irrigation Regimes. Front. Plant Sci. 12, 1–11. doi: 10.3389/fpls.2021.657578.

Liang, T., Duan, B., Luo, X., Ma, Y., and Yuan, Z. (2021). Identification of High Nitrogen Use Efficiency Phenotype in Rice (Oryza sativa L.) Through Entire Growth Duration by Unmanned Aerial Vehicle Multispectral Imagery. 12. doi: 10.3389/fpls.2021.740414.

Liu, H., Zhu, H., and Wang, P. (2017). Quantitative modelling for leaf nitrogen content of winter wheat using UAV-based hyperspectral data. Int. J. Remote Sens. 38, 2117–2134. doi: 10.1080/01431161.2016.1253899.

Liu, S., Li, L., Gao, W., Zhang, Y., Liu, Y., Wang, S., et al. (2018). Diagnosis of nitrogen status in winter oilseed rape (Brassica napus L.) using in-situ hyperspectral data and unmanned aerial vehicle (UAV) multispectral images. Comput. Electron. Agric. 151, 185–195. doi: 10.1016/j.compag.2018.05.026.

Lu, N., Wang, W., Zhang, Q., Li, D., Yao, X., Tian, Y., et al. (2019). Estimation of Nitrogen Nutrition Status in Winter Wheat From Unmanned Aerial Vehicle Based Multi-Angular Multispectral Imagery. Front. Plant Sci. 10. doi: 10.3389/fpls.2019.01601.

Maresma, Á., Ariza, M., Martníez, E., Lloveras, J., and Martínez-Casasnovas, J. A. (2016). Analysis of vegetation indices to determine nitrogen application and yield prediction in maize (zea mays l.) from a standard uav service. Remote Sens. 8. doi: 10.3390/rs8120973.

Maydup, M. L., Antonietta, M., Guiamet, J. J., and Tambussi, E. A. (2012). The contribution of green parts of the ear to grain filling in old and modern cultivars of bread wheat (Triticum aestivum L.): Evidence for genetic gains over the past century. F. Crop. Res. 134, 208–215. doi: 10.1016/j.fcr.2012.06.008.

Mevik, B. H., and Wehrens, R. (2007). The pls package: Principal component and partial least squares regression in R. J. Stat. Softw. 18, 1–23. doi: 10.18637/jss.v018.i02.

Murray, H., Lucieer, A., and Williams, R. (2010). Texture-based classification of sub-Antarctic vegetation communities on Heard Island. Int. J. Appl. Earth Obs. Geoinf. 12, 138–149. doi: 10.1016/j.jag.2010.01.006.

Nigon, T. J., Mulla, D. J., Rosen, C. J., Cohen, Y., Alchanatis, V., Knight, J., et al. (2015). Hyperspectral aerial imagery for detecting nitrogen stress in two potato cultivars. Comput. Electron. Agric. 112, 36–46. doi: 10.1016/j.compag.2014.12.018.

Nigon, T. J., Yang, C., Paiao, G. D., Mulla, D. J., Knight, J. F., and Fernández, F. G. (2020). Prediction of early season nitrogen uptake in maize using high-resolution aerial hyperspectral imagery. Remote Sens. 12. doi: 10.3390/RS12081234.

Ohyama Takuji (2010). Nitrogen as a major essential element of plants. Nitrogen Assim. plants 37, 2– 17.

Onojeghuo, A. O., Blackburn, G. A., Huang, J., Kindred, D., and Huang, W. (2018). Applications of satellite ‘hyper-sensing’ in Chinese agriculture: Challenges and opportunities. Int. J. Appl. Earth Obs. Geoinf. 64, 62–86. doi: 10.1016/j.jag.2017.09.005.

Sanchez-Bragado, R., Elazab, A., Zhou, B., Serret, M. D., Bort, J., Nieto-Taladriz, M. T., et al. (2014). Contribution of the ear and the flag leaf to grain filling in durum wheat inferred from the carbon isotope signature: Genotypic and growing conditions effects. J. Integr. Plant Biol. 56, 444–454. doi: 10.1111/jipb.12106.

Sanchez-Bragado, R., Molero, G., Reynolds, M. P., and Araus, J. L. (2016). Photosynthetic contribution of the ear to grain filling in wheat: A comparison of different methodologies for evaluation. J. Exp. Bot. 67, 2787–2798. doi: 10.1093/jxb/erw116.

Schroder, J. L., Zhang, H., Girma, K., Raun, W. R., Penn, C. J., and Payton, M. E. (2011). Soil Acidification from Long-Term Use of Nitrogen Fertilizers on Winter Wheat. Soil Sci. Soc. Am. J. 75, 957–964. doi: 10.2136/sssaj2010.0187.

Sinclair, T. R., Rufty, T. W., and Lewis, R. S. (2019). Increasing Photosynthesis: Unlikely Solution For World Food Problem. Trends Plant Sci. 24, 1032–1039. doi: 10.1016/j.tplants.2019.07.008.

Song, X., Feng, W., He, L., Xu, D., Zhang, H. Y., Li, X., et al. (2016). Examining view angle effects on leaf N estimation in wheat using field reflectance spectroscopy. ISPRS J. Photogramm. Remote Sens. 122, 57–67. doi: 10.1016/j.isprsjprs.2016.10.002.

Stroppiana, D., Boschetti, M., Brivio, P. A., and Bocchi, S. (2009). Plant nitrogen concentration in paddy rice from field canopy hyperspectral radiometry. F. Crop. Res. 111, 119–129. doi: 10.1016/j.fcr.2008.11.004.

T., T., I., A., and T., O. (1986). The diagnosis of nitrogen nutrition of rice plants (Sasanishiki) using chlorophyll-meter. Japanese J. Soil Sci. Plant Nutr. v. 57.

Tian, Y. C., Yao, X., Yang, J., Cao, W. X., Hannaway, D. B., and Zhu, Y. (2011). Assessing newly developed and published vegetation indices for estimating rice leaf nitrogen concentration with ground- and space-based hyperspectral reflectance. F. Crop. Res. 120, 299–310. doi: 10.1016/j.fcr.2010.11.002.

Tilly, N., and Bareth, G. (2019). Estimating nitrogen from structural crop traits at field scale-a novel approach versus spectral vegetation indices. Remote Sens. 11. doi: 10.3390/rs11172066.

Vergara-Diaz, O., Vatter, T., Kefauver, S. C., Obata, T., Fernie, A. R., and Araus, J. L. (2020). Assessing durum wheat ear and leaf metabolomes in the field through hyperspectral data. Plant J. 102, 615–630. doi: 10.1111/tpj.14636.

Verrelst, J., Camps-Valls, G., Muñoz-Marí, J., Rivera, J. P., Veroustraete, F., Clevers, J. G. P. W., et al. (2015). Optical remote sensing and the retrieval of terrestrial vegetation bio-geophysical properties - A review. ISPRS J. Photogramm. Remote Sens. 108, 273–290. doi: 10.1016/j.isprsjprs.2015.05.005.

Vicente, R., Vergara-Díaz, O., Medina, S., Chairi, F., Kefauver, S. C., Bort, J., et al. (2018). Durum wheat ears perform better than the flag leaves under water stress: Gene expression and physiological evidence. Environ. Exp. Bot. 153, 271–285. doi: 10.1016/j.envexpbot.2018.06.004.

Wan, S., and Chang, S. H. (2019). Crop classification with WorldView-2 imagery using Support Vector Machine comparing texture analysis approaches and grey relational analysis in Jianan Plain, Taiwan. Int. J. Remote Sens. 40, 8076–8092. doi: 10.1080/01431161.2018.1539275.

Wang, F., Yang, M., Ma, L., Zhang, T., Qin, W., Li, W., et al. (2022a). Estimation of Aboveground Biomass of Winter Wheat Based on Consumer-Grade Multi-Spectral UAV. 14, 1251. Available at: https://doi.org/10.3390/rs14051251.

Wang, L., Chang, Q., Li, F., Yan, L., Huang, Y., Wang, Q., et al. (2019). Effects of growth stage development on paddy rice leaf area index prediction models. Remote Sens. 11, 1–18. doi: 10.3390/rs11030361.

Wang, W., Yao, X., Yao, X. F., Tian, Y. C., Liu, X. J., Ni, J., et al. (2012). Estimating leaf nitrogen concentration with three-band vegetation indices in rice and wheat. F. Crop. Res. 129, 90–98. doi: 10.1016/j.fcr.2012.01.014.

Wang, Y. P., Chang, Y. C., and Shen, Y. (2022b). Estimation of nitrogen status of paddy rice at vegetative phase using unmanned aerial vehicle based multispectral imagery. Precis. Agric. 23. doi: 10.1007/s11119-021-09823-w.

Wang, Z., Skidmore, A. K., Darvishzadeh, R., Heiden, U., Heurich, M., and Wang, T. (2015). Leaf Nitrogen Content Indirectly Estimated by Leaf Traits Derived from the PROSPECT Model. IEEE J. Sel. Top. Appl. Earth Obs. Remote Sens. 8, 3172–3182. doi: 10.1109/JSTARS.2015.2422734.

Wang, Z., Zhang, W., Beebout, S. S., Zhang, H., Liu, L., Yang, J., et al. (2016). Grain yield, water and nitrogen use efficiencies of rice as influenced by irrigation regimes and their interaction with nitrogen rates. F. Crop. Res. 193, 54–69. doi: 10.1016/j.fcr.2016.03.006.

Wen, P., Shi, Z., Li, A., Ning, F., Zhang, Y., Wang, R., et al. (2021). Estimation of the vertically integrated leaf nitrogen content in maize using canopy hyperspectral red edge parameters. Precis. Agric. 22, 984–1005. doi: 10.1007/s11119-020-09769-5.

Wold, S., Sjöström, M., and Eriksson, L. (2001). PLS -regression: A basic tool of chemometrics. Chemom. Intell. Lab. Syst. 58, 109–130. doi: 10.1016/S0169-7439(01)00155-1.

Xue, L., Cao, W., Luo, W., Dai, T., and Zhu, Y. (2004). Monitoring Leaf Nitrogen Status in Rice with Canopy Spectral Reflectance. Agron. J. 96, 135–142. doi: 10.2134/agronj2004.0135.

Yang, G., Zhao, C., Pu, R., Feng, H., Li, Z., Li, H., et al. (2015). Leaf nitrogen spectral reflectance model of winter wheat (Triticum aestivum) based on PROSPECT: simulation and inversion. J. Appl. Remote Sens. 9, 095976. doi: 10.1117/1.jrs.9.095976.

Yang, K., Gong, Y., Fang, S., Duan, B., Yuan, N., Peng, Y., et al. (2021). Combining spectral and texture features of uav images for the remote estimation of rice lai throughout the entire growing season. Remote Sens. 13. doi: 10.3390/rs13153001.

Yang, M., Hassan, M. A., Xu, K., Zheng, C., Rasheed, A., Zhang, Y., et al. (2020). Assessment of Water and Nitrogen Use Efficiencies Through UAV-Based Multispectral Phenotyping in Winter Wheat. Front. Plant Sci. 11, 1–16. doi: 10.3389/fpls.2020.00927.

Yao, X., Huang, Y., Shang, G., Zhou, C., Cheng, T., Tian, Y., et al. (2015). Evaluation of six algorithms to monitor wheat leaf nitrogen concentration. Remote Sens. 7, 14939–14966. doi: 10.3390/rs71114939.

Yu, K., Lenz-Wiedemann, V., Chen, X., and Bareth, G. (2014). Estimating leaf chlorophyll of barley at different growth stages using spectral indices to reduce soil background and canopy structure effects. ISPRS J. Photogramm. Remote Sens. 97, 58–77. doi: 10.1016/j.isprsjprs.2014.08.005.

Yu, K., Li, F., Gnyp, M. L., Miao, Y., Bareth, G., and Chen, X. (2013). Remotely detecting canopy nitrogen concentration and uptake of paddy rice in the Northeast China Plain. ISPRS J. Photogramm. Remote Sens. 78, 102–115. doi: 10.1016/j.isprsjprs.2013.01.008.

Yuan, Z., Ata-Ul-Karim, S. T., Cao, Q., Lu, Z., Cao, W., Zhu, Y., et al. (2016). Indicators for diagnosing nitrogen status of rice based on chlorophyll meter readings. F. Crop. Res. 185, 12–20. doi: 10.1016/j.fcr.2015.10.003.

Yue, J., Yang, G., Tian, Q., Feng, H., Xu, K., and Zhou, C. (2019). Estimate of winter-wheat aboveground biomass based on UAV ultrahigh-ground-resolution image textures and vegetation indices. ISPRS J. Photogramm. Remote Sens. 150, 226–244. doi: 10.1016/j.isprsjprs.2019.02.022.

Zhang, H. Y., Ren, X. X., Zhou, Y., Wu, Y. P., He, L., Heng, Y. R., et al. (2018). Remotely assessing photosynthetic nitrogen use efficiency with in situ hyperspectral remote sensing in winter wheat. Eur. J. Agron. 101, 90–100. doi: 10.1016/j.eja.2018.08.010.

Zheng, H., Cheng, T., Zhou, M., Li, D., Yao, X., Tian, Y., et al. (2019). Improved estimation of rice aboveground biomass combining textural and spectral analysis of UAV imagery. Precis. Agric. 20, 611–629. doi: 10.1007/s11119-018-9600-7.

Zheng, H., Li, W., Jiang, J., Liu, Y., Cheng, T., Tian, Y., et al. (2018). A comparative assessment of different modeling algorithms for estimating leaf nitrogen content in winter wheat using multispectral images from an unmanned aerial vehicle. Remote Sens. 10. doi: 10.3390/rs10122026.

Zheng, H., Ma, J., Zhou, M., Li, D., Yao, X., Cao, W., et al. (2020). Enhancing the nitrogen signals of rice canopies across critical growth stages through the integration of textural and spectral information from unmanned aerial vehicle (UAV) multispectral imagery. Remote Sens. 12. doi: 10.3390/rs12060957.

Zhu, W., Rezaei, E. E., Nouri, H., Sun, Z., Li, J., Yu, D., et al. (2022). UAV-based indicators of crop growth are robust for distinct water and nutrient management but vary between crop development phases. F. Crop. Res. 284, 108582. doi: 10.1016/j.fcr.2022.108582.

