## Supplementary Material S1-3 for "Characterization of N distribution in different organs of winter wheat using UAV-based remote sensing": Supplementary Material S1.docx

**Table S1**. Vegetation indices used in the study

| abbreviations | Vegetation index | Formula | Authors |
| --- | --- | --- | --- |
| VI.1 | BNDVI | $(NIR-BLUE)/(NIR+BLUE)$ | (Wang et al., 2007) |
| VI.2 | CI-GREEN | $(NIR/GREEN)-1$ | (Gitelson et al., 2003) |
| VI.3 | CI-RED | $(NIR/RED)-1$ | (Clevers et al., 2017) |
| VI.4 | CI-REG | $(NIR/REDEDGE)-1$ | (Gitelson et al., 2003) |
| VI.5 | CVI | $(NIR/GREEN)\times(RED/GREEN)$ | (Vincini et al., 2008) |
| VI.6 | DVI | $NIR-RED$ | (Rouse et al., 1973) |
| VI.7 | DVI-GREEN | $NIR-GREEN$ | (Rouse et al., 1973) |
| VI.8 | DVI-REG | $NIR-REDEDGE$ | (Rouse et al., 1973) |
| VI.9 | EVI | $2.5(NIR-RED)/(1+NIR-2.4RED)$ | (Huete A. et al., 2002) |
| VI.10 | EVI2 | $2.5(NIR-RED)/(NIR+2.4RED+1)$ | (Jiang et al., 2008) |
| VI.11 | GARI | $\frac{NIR-[GREEN-1.7\left( BLUE-RED \right)]}{NIR+[GREEN-1.7\left( BLUE-RED \right)]}$ | (Gitelson et al., 1996) |
| VI.12 | GNDVI | $(NIR-GREEN)/(NIR+GREEN)$ | (Gitelson et al., 2003) |
| VI.13 | GOSAVI | $(NIR-GREEN)/(NIR+GREEN+0.16)$ | (Siegmann et al., 2014) |
| VI.14 | GRVI | $(GREEN-RED)/(GREEN+RED)$ | (Tucker, 1979) |
| VI.15 | LCI | $(NIR-REDEDGE)/(NIR-RED)$ | (Xiao et al., 2014) |
| VI.16 | MCARI | $\left[ \left( REDEDGE-RED \right)-0.2\left( REDEDGE-GREEN \right) \right]*(REDEDGE/RED)$ | (Daughtry et al., 2000) |
| VI.17 | MCARI1 | $1.2[2.5\left( NIR-RED \right)-1.3\left( NIR-GREEN \right)]$ | (Haboudane et al., 2004) |
| VI.18 | MCARI2 | $\frac{3.75\left( NIR-RED \right)-1.95(NIR-GREEN)}{\sqrt{{(2NIR+1)}^{2}-6\left( NIR-5\sqrt{RED} \right)-0.5}}$ | (Haboudane et al., 2004) |
| VI.19 | MNLI | $(1.5{NIR}^{2}-1.5GREEN)/({NIR}^{2}+RED+0.5)$ | (Gong et al., 2003) |
| VI.20 | MSR | $[(NIR/RED)-1]/\surd((NIR/RED)+1)$ | (Chen, 1996) |
| VI.21 | MSR-REG | $[(NIR/REDEDGE)-1]/\surd((NIR/REDEDGE)+1)$ | (Chen, 1996) |
| VI.22 | MTCI | $(NIR-REG)/(NIR-RED)$ | (Dash and Curran, 2004) |
| VI.23 | NDRE | $(NIR-REDEDGE)/(NIR+REDEDGE)$ | (Gitelson and Merzlyak, 1997) |
| VI.24 | NDREI | $(REDEDGE-GREEN)/(REDEDGE+GREEN)$ | (Hassan et al., 2018) |
| VI.25 | NAVI | $1-RED/NIR$ | (Agapiou et al., 2013) |
| VI.26 | NDVI | $(NIR-RED)/(NIR+RED)$ | (Rouse et al., 1973) |
| VI.27 | OSAVI | $1.6[(NIR-RED)/(NIR+RED+0.16)]$ | (Rondeaux et al., 1996) |
| VI.28 | OSAVI-REG | $1.6[(NIR-REDEDGE)/(NIR+REDEDGE+0.16)]$ | (Rondeaux et al., 1996) |
| VI.29 | RDVI | $(NIR-RED)/\sqrt{(NIR+RED)}$ | (Roujean and Breon, 1995) |
| VI.30 | RDVI-REG | $(NIR-REDEDGE)/\sqrt{(NIR+REDEDGE)}$ | (Roujean and Breon, 1995) |
| VI.31 | RGBVI | $({GREEN}^{2}-BLUE*RED)/({GREEN}^{2}+BLUE*RED)$ | (Bendig et al., 2015) |
| VI.32 | RTVI-CORE | $100\left( NIR-REDDEGE \right)-10(NIR-GREEN)$ | (Walsh et al., 2018) |
| VI.33 | RVI | $NIR/RED$ | (Rouse et al., 1973) |
| VI.34 | SAVI | $1.5(NIR-RED)/(NIR+RED+0.5)$ | (Huete, 1988) |
| VI.35 | SAVI-GREEN | $1.5(NIR-GREEN)/(NIR+GREEN+0.5)$ | (Verrelst et al., 2008) |
| VI.36 | S-CCCI | $NDRE/NDVI$ | (Raper and Varco, 2015) |
| VI.37 | SIPI | $(NIR-BLUE)/(NIR-RED)$ | (Peñuelas et al., 1994) |
| VI.38 | SR-REG | $NIR/REDEDGE$ | (Walsh et al., 2018) |
| VI.39 | TCARI | $3[(REDEDGE-RED)-0.2(REDEDGE-GREEN)*(REDEDGE/RED)]$ | (Haboudane et al., 2002) |
| VI.40 | TCARI/OSAVI | $TCARI/OSAVI$ | (Haboudane et al., 2002) |
| VI.41 | TVI | $\left[ 120\left( NIR-GREEN \right)-200\left( RED-GREEN \right) \right]/2$ | (Broge and Leblanc, 2001) |
| VI.42 | VARI | $(GREEN-RED)/(GREEN+RED-BLUE)$ | (Gitelson, 2013) |
| VI.43 | WDRVI | $(0.2NIR-RED)/(0.2NIR+RED)$ | (Pilson and Decker, 2002) |

**Table S2**. Texture features used in this study

| abbreviations | Texture feature | abbreviations | Texture feature |
| --- | --- | --- | --- |
| TF.1 | B_con | TF.21 | Nir_ho |
| TF.2 | B_cor | TF.22 | Nir_mean |
| TF.3 | B_dis | TF.23 | Nir_se |
| TF.4 | B_en | TF.24 | Nir_var |
| TF.5 | B_ho | TF.25 | R_con |
| TF.6 | B_mean | TF.26 | R_cor |
| TF.7 | B_se | TF.27 | R_dis |
| TF.8 | B_var | TF.28 | R_en |
| TF.9 | G_con | TF.29 | R_ho |
| TF.10 | G_cor | TF.30 | R_mean |
| TF.11 | G_dis | TF.31 | R_se |
| TF.12 | G_en | TF.32 | R_var |
| TF.13 | G_ho | TF.33 | Reg_con |
| TF.14 | G_mean | TF.34 | Reg_cor |
| TF.15 | G_se | TF.35 | Reg_dis |
| TF.16 | G_var | TF.36 | Reg_en |
| TF.17 | Nir_con | TF.37 | Reg_ho |
| TF.18 | Nir_cor | TF.38 | Reg_mean |
| TF.19 | Nir_dis | TF.39 | Reg_se |
| TF.20 | Nir_en | TF.40 | Reg_var |

The following formulars are for the calculation of the texture features.

| $mean= \sum_{i} \sum_{j} p\left( i, j \right)i$ | (01) |
| --- | --- |
| $var= \sum_{i} \sum_{j} \left( i-u \right)^{2}p(i, j)$ | (02) |
| $ho= \sum_{i} \sum_{j} \frac{1}{1+\left( i-j \right)^{2}}p(i, j)$ | (03) |
| $con= \sum_{n=0}^{N_{g}-1} n^{2}\left\{ \begin{aligned} \sum_{i=1}^{N_{g}} \sum_{j=1}^{N_{g}} p(i, j) \\ \left\vert i-j \right\vert=n \end{aligned} \right\}$ | (04) |
| $dis= \sum_{n=1}^{N_{g}-1} n\left\{ \begin{aligned} \sum_{i=1}^{N_{g}} \sum_{j=1}^{N_{g}} p(i, j) \\ \left\vert i-j \right\vert=n \end{aligned} \right\}$ | (05) |
| $en= -\sum_{i} \sum_{j} p\left( i,j \right)log(p\left( i, j \right))$ | (06) |
| $se= \sum_{i} \sum_{j} {\{p(i, j)\}}^{2}$ | (07) |
| $cor= \frac{\sum_{i} \sum_{j} \left( i, j \right)p\left( i,j \right)-\mu_{x}\mu_{y}}{\sigma_{x}\sigma_{y}}$ | (08) |

Where, p(i, j)*p(i, j)* denotes the (i, j)*(i, j)*th entry in a normalized GLCM $P(i,j)/Rp_{x}(i)[=P(i,j)/R$; *p_x_*(*i*) denotes the *ii* th entry in the marginal-probability matrix which was calculated via summing the rows of $p(i,j)\sum_{j=1}^{N_{g}} P\left( i,j \right)N_{g}p\left( i,j \right)=\sum_{j=1}^{N_{g}} P\left( i,j \right);$ N_g_ denotes the number of distinct grey levels in the UAV quantized imagery; p_y_(j) equals $\sum_{i=1}^{N_{g}} p\left( i,j \right)p_{y}(j)$, i.e., $\sum_{i=1}^{N_{g}} p(i, j)$; *μ_x_*, *μ_y_*, *σ_x_*, and *σ_y_* denote the mean and standard deviation values of *p_x_* and *p_y_*, respectively.

**Table S3**. Descriptive statistics of nitrogen content and biomass in the vegetative growth phase and reproductive growth phase of winter wheat under different N levels.

| Growth phase | n | Min | Max | Mean | SD | CV (%) |
| --- | --- | --- | --- | --- | --- | --- |
| Leaf NC(%)  Vegetative growth (GS 31-50)  Reproductive growth (GS 70-90)  All pooled data | 45  75  120 | 2.24  0.91  0.91 | 4.95  3.64  4.95 | 3.68  2.27  2.80 | 0.62  0.76  0.98 | 16.84  33.41  35.20 |
| Leaf DMW(t/ha)  Vegetative growth (GS 31-50)  Reproductive growth (GS 70-90)  All pooled data | 45  75  120 | 1.22  0.84  0.84 | 4.18  3.93  4.18 | 2.95  2.56  5.71 | 9.16  7.88  8.56 | 31.01  30.76  31.61 |
| Stem NC(%)  Vegetative growth (GS 31-50)  Reproductive growth (GS 70-90)  All pooled data | 45  75  120 | 0.85  0.29  0.29 | 1.81  1.31  1.81 | 1.38  0.79  1.01 | 0..27  0.25  0.38 | 19.48  31.10  37.73 |
| Stem DMW(t/ha)  Vegetative growth (GS 31-50)  Reproductive growth (GS 70-90)  All pooled data | 45  75  120 | 2.72  5.14  2.72 | 9.87  13.80  13.80 | 6.50  9.64  8.46 | 1.93  2.26  2.62 | 29.71  23.44  31.00 |
| Spike NC(%)  Vegetative growth (GS 31-50)  Reproductive growth (GS 70-90)  All pooled data | 30  75  105 | 1.99  1.41  1.41 | 4.73  2.60  4.73 | 2.90  1.88  2.17 | 0.90  0.22  0.69 | 31.01  12.00  31.60 |
| Spike DMW(t/ha)  Vegetative growth (GS 31-50)  Reproductive growth (GS 70-90)  All pooled data | 30  75  105 | 0.42  1.42  0.42 | 2.37  11.20  11.20 | 1.29  5.97  4.63 | 0.66  2.86  3.23 | 51.53  47.86  69.75 |
| Grain NC(%)  Vegetative growth (GS 31-50)  Reproductive growth (GS 70-90)  All pooled data | -  75  75 | -  1.62  1.62 | -  3.06  3.06 | -  2.41  2.41 | -  0.36  0.36 | -  15.04  15.04 |
| Grain DMW(t/ha)  Vegetative growth (GS 31-50)  Reproductive growth (GS 70-90)  All pooled data | -  75  75 | -  0.26  0.26 | -  8.14  8.14 | -  3.47  3.47 | -  2.58  2.58 | -  74.41  74.41 |
| Plant NC(%)  Vegetative growth (GS 31-50)  Reproductive growth (GS 70-90)  All pooled data | 45  75  120 | 1.45  0.80  0.80 | 3.00  1.85  3.00 | 2.16  1.37  1.67 | 0.45  0.24  0.51 | 20.86  17.78  30.74 |
| Plant DMW(t/ha)  Vegetative growth (GS 31-50)  Reproductive growth (GS 70-90)  All pooled data | 45  75  120 | 3.97  8.26  3.97 | 15.96  27.08  27.08 | 10.31  18.16  15.22 | 3.27  4.37  5.51 | 31.71  24.06  36.22 |

**Table S4**. The top 5 most relevant image features with NC of organs and the whole plant.

| Data set | Part of winter wheat | Vegetative growth phase | Reproductive growth phase |
| --- | --- | --- | --- |
| VIs |  |  |  |
|  | Leaf | RGBVI, GRVI, MTCI, SIPI, NDREI | GOSAVI, TVI, MSR, RTVI-CORE, RDVI |
|  | Stem | MCARI, GRVI, MSR, NDREI, WDRVI | MSR-REG, NDRE, RDVI-REG, CVI, CI-REG |
|  | Spike | MCARI2, LCI, CI-RED, RVI, S-CCCI | DVI-REG, CVI, MSR-REG, SAVI, RDVI-CORE |
|  | Grain | - | RTVI-CORE, RDVI-REG, DVI-REG, TVI, CI-REG |
|  | Plant | RGBVI, GRVI, MCARI, NDREI, WDRVI | CVI, MSR-REG, DVI-REG, RDVI-REG, NDRE |
| TFs |  |  |  |
|  | Leaf | Reg_mean, B_cor, B_mean, G_cor, G_mean | R_ho, R_dis, R_con G_en, R_en |
|  | Stem | G_cor, G_mean, R_mean, B_mean, REG_mean | G_mean, R_mean, B_mean, Reg_mean, R_cor |
|  | Spike | R_con, R_dis, R_en, R_ho, R_se | Reg_mean, B_cor, G_mean, B_mean, R_cor |
|  | Grain | - | G_mean, R_mean, B_mean, R_ho, R_dis |
|  | Plant | Reg_mean, G_cor, B_mean, B_cor, G_mean | Reg_mean, G_mean, B_mean, R_cor, B_cor |

**Table S5**. Number of image features selected by the PLSR and RF models in different growth phases.

| Data set | Part of winter wheat | Vegetative growth | | Reproductive growth | |
| --- | --- | --- | --- | --- | --- |
|  |  | PLSR | RF | PLSR | RF |
| VIs |  |  |  |  |  |
|  | Leaf | 10 | 11 | 29 | 26 |
|  | Stem | 19 | 20 | 26 | 18 |
|  | Spike | 3 | 6 | 13 | 8 |
|  | Grain | - | - | 9 | 18 |
|  | Plant | 10 | 13 | 13 | 12 |
| TFs |  |  | |  | |
|  | Leaf | 15 | 11 | 17 | 14 |
|  | Stem | 14 | 8 | 18 | 12 |
|  | Spike | 20 | 17 | 14 | 7 |
|  | Grain | - | - | 19 | 13 |
|  | Plant | 14 | 7 | 15 | 7 |
| VIs + TFs |  |  | |  | |
|  | Leaf | 34 | 17 | 44 | 41 |
|  | Stem | 49 | 15 | 46 | 20 |
|  | Spike | 26 | 27 | 34 | 11 |
|  | Grain | - | - | 32 | 26 |
|  | Plant | 39 | 12 | 35 | 17 |

**References**

Agapiou, A., Alexakis, D. D., Stavrou, M., Sarris, A., Themistocleous, K., and Hadjimitsis, D. G. (2013). Prospects and limitations of vegetation indices in archeological research: the Neolithic Thessaly case study. *Earth Resour. Environ. Remote Sensing/GIS Appl. IV* 8893, 88930D. doi: 10.1117/12.2028661.

Bendig, J., Yu, K., Aasen, H., Bolten, A., Bennertz, S., Broscheit, J., et al. (2015). Combining UAV-based plant height from crop surface models, visible, and near infrared vegetation indices for biomass monitoring in barley. *Int. J. Appl. Earth Obs. Geoinf.* 39, 79–87. doi: 10.1016/j.jag.2015.02.012.

Broge, N. H., and Leblanc, E. (2001). Comparing prediction power and stability of broadband and hyperspectral vegetation indices for estimation of green leaf area index and canopy chlorophyll density. *Remote Sens. Environ.* 76, 156–172. doi: 10.1016/S0034-4257(00)00197-8.

Chen, J. M. (1996). Evaluation of vegetation indices and a modified simple ratio for boreal applications. *Can. J. Remote Sens.* 22, 229–242. doi: 10.1080/07038992.1996.10855178.

Clevers, J. G. P. W., Kooistra, L., and van den Brande, M. M. M. (2017). Using Sentinel-2 data for retrieving LAI and leaf and canopy chlorophyll content of a potato crop. *Remote Sens.* 9, 1–15. doi: 10.3390/rs9050405.

Dash, J., and Curran, P. J. (2004). The MERIS terrestrial chlorophyll index. *Int. J. Remote Sens.* 25, 5403–5413. doi: 10.1080/0143116042000274015.

Daughtry, C. S. T., Walthall, C. L., Kim, M. S., De Colstoun, E. B., and McMurtrey, J. E. (2000). Estimating corn leaf chlorophyll concentration from leaf and canopy reflectance. *Remote Sens. Environ.* 74, 229–239. doi: 10.1016/S0034-4257(00)00113-9.

Gitelson, A. A. (2013). Remote estimation of crop fractional vegetation cover: The use of noise equivalent as an indicator of performance of vegetation indices. *Int. J. Remote Sens.* 34, 6054–6066. doi: 10.1080/01431161.2013.793868.

Gitelson, A. A., Gritz, Y., and Merzlyak, M. N. (2003). Relationships between leaf chlorophyll content and spectral reflectance and algorithms for non-destructive chlorophyll assessment in higher plant leaves. *J. Plant Physiol.* 160, 271–282. doi: 10.1078/0176-1617-00887.

Gitelson, A. A., and Merzlyak, M. N. (1997). Remote estimation of chlorophyll content in higher plant leaves. *Int. J. Remote Sens.* 18, 2691–2697. doi: 10.1080/014311697217558.

Gitelson, A. A., Merzlyak, M. N., and Lichtenthaler, H. K. (1996). Detection of red edge position and chlorophyll content by reflectance measurements near 700 nm. *J. Plant Physiol.* 148, 501–508. doi: 10.1016/S0176-1617(96)80285-9.

Gong, P., Pu, R., Biging, G. S., and Larrieu, M. R. (2003). Estimation of forest leaf area index using vegetation indices derived from Hyperion hyperspectral data. *IEEE Trans. Geosci. Remote Sens.* 41, 1355–1362. doi: 10.1109/TGRS.2003.812910.

Haboudane, D., Miller, J. R., Pattey, E., Zarco-Tejada, P. J., and Strachan, I. B. (2004). Hyperspectral vegetation indices and novel algorithms for predicting green LAI of crop canopies: Modeling and validation in the context of precision agriculture. *Remote Sens. Environ.* 90, 337–352. doi: 10.1016/j.rse.2003.12.013.

Haboudane, D., Miller, J. R., Tremblay, N., Zarco-Tejada, P. J., and Dextraze, L. (2002). Integrated narrow-band vegetation indices for prediction of crop chlorophyll content for application to precision agriculture. *Remote Sens. Environ.* 81, 416–426. doi: 10.1016/S0034-4257(02)00018-4.

Hassan, M. A., Yang, M., Rasheed, A., Jin, X., Xia, X., Xiao, Y., et al. (2018). Time-series multispectral indices from unmanned aerial vehicle imagery reveal senescence rate in bread wheat. *Remote Sens.* 10. doi: 10.3390/rs10060809.

Huete A., Didan, K., T. Miura, E.P. Rodriguez a, X. Gao, and L.G. Ferreira (2002). Overview of the radiometric and biophysical performance of the MODIS vegetation indices. *Remote Sens. Environ.* 83, 195–213. doi: 10.1016/S0034-4257(02)00096-2.

Huete, A. R. (1988). A soil-adjusted vegetation index (SAVI). *Remote Sens. Environ.* 25, 295–309. doi: 10.1016/0034-4257(88)90106-X.

Jiang, Z., Huete, A. R., Didan, K., and Miura, T. (2008). Development of a two-band enhanced vegetation index without a blue band. *Remote Sens. Environ.* 112, 3833–3845. doi: 10.1016/j.rse.2008.06.006.

Peñuelas, J., Gamon, J. A., Fredeen, A. L., Merino, J., and Field, C. B. (1994). Reflectance indices associated with physiological changes in nitrogen- and water-limited sunflower leaves. *Remote Sens. Environ.* 48, 135–146. doi: 10.1016/0034-4257(94)90136-8.

Pilson, D., and Decker, K. L. (2002). Compensation for herbivory in wild sunflower: Response to simulated damage by the head-clipping weevil. *Ecology* 83, 3097–3107. doi: 10.1890/0012-9658(2002)083[3097:CFHIWS]2.0.CO;2.

Raper, T. B., and Varco, J. J. (2015). Canopy-scale wavelength and vegetative index sensitivities to cotton growth parameters and nitrogen status. *Precis. Agric.* 16, 62–76. doi: 10.1007/s11119-014-9383-4.

Rondeaux, G., Steven, M., and Baret, F. (1996). Optimization of soil-adjusted vegetation indices. *Remote Sens. Environ.* 55, 95–107. doi: 10.1016/0034-4257(95)00186-7.

Roujean, J. L., and Breon, F. M. (1995). Estimating PAR absorbed by vegetation from bidirectional reflectance measurements. *Remote Sens. Environ.* 51, 375–384. doi: 10.1016/0034-4257(94)00114-3.

Rouse, J. W., Haas, R. H., Schell, J. A., and Deeering, D. . (1973). Monitoring vegetation systems in the Great Plains with ERTS (Earth Resources Technology Satellite). in *Third Earth Resources Technology Satellite-1 Symposium*, 309–317.

Siegmann, B., Jarmer, T., Lilienthal, H., Richter, N., Selige, T., and Höfled, B. (2014). Comparison of narrow band vegetation indices and empirical models from hyperspectral remote sensing data for the assessment of wheat nitrogen concentration. *ISPRS Work. UAV-based Remote Sens. Methods Monit. Veg.*, 1–2.

Tucker, C. J. (1979). Red and photographic infrared linear combinations for monitoring vegetation. *Remote Sens. Environ.* 8, 127–150. doi: 10.1016/0034-4257(79)90013-0.

Verrelst, J., Schaepman, M. E., Koetz, B., and Kneubühler, M. (2008). Angular sensitivity analysis of vegetation indices derived from CHRIS/PROBA data. *Remote Sens. Environ.* 112, 2341–2353. doi: 10.1016/j.rse.2007.11.001.

Vincini, M., Frazzi, E., and D’Alessio, P. (2008). A broad-band leaf chlorophyll vegetation index at the canopy scale. *Precis. Agric.* 9, 303–319. doi: 10.1007/s11119-008-9075-z.

Walsh, O. S., Shafian, S., Marshall, J. M., Jackson, C., McClintick-Chess, J. R., Blanscet, S. M., et al. (2018). Assessment of UAV Based Vegetation Indices for Nitrogen Concentration Estimation in Spring Wheat. *Adv. Remote Sens.* 07, 71–90. doi: 10.4236/ars.2018.72006.

Wang, F., Huang, J., Tang, Y., and Wang, X. (2007). New Vegetation Index and Its Application in Estimating Leaf Area Index of Rice. *Rice Sci.* 14, 195–203. doi: 10.1016/s1672-6308(07)60027-4.

Xiao, Y., Zhao, W., Zhou, D., and Gong, H. (2014). Sensitivity analysis of vegetation reflectance to biochemical and biophysical variables at leaf, canopy, and regional scales. *IEEE Trans. Geosci. Remote Sens.* 52, 4014–4024. doi: 10.1109/TGRS.2013.2278838.
